# Native KCC2 interactome reveals PACSIN1 as a critical regulator of synaptic inhibition

**DOI:** 10.1101/142265

**Authors:** Vivek Mahadevan, C. Sahara Khademullah, Zahra Dargaei, Jonah Chevrier, Pavel Uvarov, Julian Kwan, Richard D. Bagshaw, Tony Pawson, Andrew Emili, Yves DeKoninck, Victor Anggono, Matti S. Airaksinen, Melanie A. Woodin

## Abstract

KCC2 is a neuron-specific K^+^-Cl^−^ cotransporter essential for establishing the Cl^−^ gradient required for hyperpolarizing inhibition. KCC2 is highly localized to excitatory synapses where it regulates spine morphogenesis and AMPA receptor confinement. Aberrant KCC2 function contributes to numerous human neurological disorders including epilepsy and neuropathic pain. Using unbiased functional proteomics, we identified the KCC2-interactome in the mouse brain to determine KCC2-protein interactions that regulate KCC2 function. Our analysis revealed that KCC2 interacts with a diverse set of proteins, and its most predominant interactors play important roles in postsynaptic receptor recycling. The most abundant KCC2 interactor is a neuronal endocytic regulatory protein termed PACSIN1 (SYNDAPIN1). We verified the PACSIN1-KCC2 interaction biochemically and demonstrated that shRNA knockdown of PACSIN1 in hippocampal neurons significantly increases KCC2 expression and hyperpolarizes the reversal potential for Cl^−^. Overall, our global native-KCC2 interactome and subsequent characterization revealed PACSIN1 as a novel and potent negative regulator of KCC2.

## Introduction

GABA and glycine are the key inhibitory neurotransmitters of the mature nervous system, and most synaptic inhibition is mediated by Cl^−^ permeable GABA_A_ and glycine receptors. This hyperpolarizing inhibition results from the inward gradient for Cl^−^ established primarily by the K^+^-Cl^−^ cotransporter KCC2, which exports Cl^−^ to maintain low intracellular Cl^−^ (Rivera et al., 1999; Doyon et al., 2016). KCC2 is a member of the *SLC12A* family of cation-chloride cotransporters and is unique among the members because it is present exclusively in neurons, and mediates the electroneutral outward cotransport of K^+^ and Cl^−^.

During embryonic development KCC2 expression is low and GABA and glycine act as excitatory neurotransmitters, however during early postnatal development KCC2 expression is dramatically upregulated and GABA and glycine become inhibitory (Ben-Ari, 2002; Blaesse et al., 2009). Excitation-inhibition imbalance underlies numerous neurological disorders (Kahle et al., 2008; Nelson and Valakh, 2015), and in many of these disorders, the decrease in inhibition results from a reduction in KCC2 expression. In particular, KCC2 dysfunction contributes to the onset of seizures (Huberfeld et al., 2007; Kahle et al., 2014; Puskarjov et al., 2014; Stödberg et al., 2015; Saitsu et al., 2016), neuropathic pain (Coull et al., 2003), schizophrenia (Tao et al., 2012), and autism spectrum disorders (ASD) (Cellot and Cherubini, 2014; Tang et al., 2015; Banerjee et al., 2016). Despite the critical importance of this transporter in maintaining inhibition and proper brain function, our understanding of KCC2 regulation is rudimentary.

In addition to its canonical role of Cl^−^ extrusion that regulates synaptic inhibition, KCC2 has also emerged as a key regulator of excitatory synaptic transmission. KCC2 is highly localized in the vicinity of excitatory synapses (Gulyas et al., 2001; Chamma et al., 2013) and regulates both the development of dendritic spine morphology (Li et al., 2007; Chevy et al., 2015; Llano et al., 2015) and function of AMPA-mediated glutamatergic synapses (Gauvain et al., 2011; Chevy et al., 2015; Llano et al., 2015). Thus, a dysregulation of these non-canonical KCC2 functions at excitatory synapses may also contribute to the onset of neurological disorders associated with both KCC2 dysfunction and excitation-inhibition imbalances.

KCC2 is regulated by multiple posttranslational mechanisms including phosphoregulation by distinct kinases and phosphatases (Lee et al., 2007; Kahle et al., 2013; Medina et al., 2014), lipid rafts and oligomerization (Blaesse et al., 2006; Watanabe et al., 2009), and protease-dependent cleavage (Puskarjov et al., 2012). KCC2 expression and function is also regulated by protein interactions, including creatine kinase B (CKB) (Inoue et al., 2006), sodium/potassium ATPase subunit 2 (ATP1A2) (Ikeda et al., 2004), chloride cotransporter interacting protein 1 (CIP1) (Wenz et al., 2009), protein associated with Myc (PAM) (Garbarini and Delpire, 2008), 4.1N (Li et al., 2007), the glutamate receptor subunit GluK2, its auxiliary subunit Neto2 (Ivakine et al., 2013; Mahadevan et al., 2014), cofilin1 (CFL1) (Chevy et al., 2015; Llano et al., 2015) and RAB11(Roussa et al., 2016). However, since KCC2 exists in a large multi-protein complex (MPC) (Mahadevan et al., 2015), it is likely that these previously identified interactions do not represent all of the components of native-KCC2 MPCs.

In the present study, we performed unbiased multi-epitope tagged affinity purifications (ME-AP) of native-KCC2 coupled with high-resolution mass spectrometry (MS) from whole-brain membrane fractions prepared from developing and mature mouse brain. We found that native KCC2 exists in macromolecular complexes comprised of interacting partners from diverse classes of transmembrane and soluble proteins. Subsequent network analysis revealed numerous previously unknown native-KCC2 protein interactors related to receptor recycling and vesicular endocytosis functions. We characterized the highest-confidence KCC2 partner identified in this screen, PACSIN1, and determined that PACSIN1 is a novel and potent negative regulator of KCC2 function.

## RESULTS

### Determining Affinity Purification (AP) conditions to extract native-KCC2

In order to determine the composition of native KCC2 MPCs using AP-MS, we first determined the detergent-based conditions that preserve native KCC2 following membrane extraction. In a non-denaturing Blue-Native PAGE (BN-PAGE), native-KCC2 migrated between 400 kDa – 1000 kDa in the presence of the native detergents C_12_E_9_, CHAPS, and DDM. However, all other detergent compositions previously used for KCC2 solubilization resulted in KCC2 migration at lower molecular weights (Figure 1a). This indicates that native detergent extractions are efficient at preserving higher-order KCC2 MPCs. Upon further analysis using standard SDS-PAGE we observed that the total KCC2 extracted was greater in C_12_E_9_ and CHAPS-based detergent extractions in comparison with all other detergents (Figure 1a, Figure 1 – Figure Supplement 1, 2), hence we restricted our further analysis to C_12_E_9_ and CHAPS-based membrane preparations.

**Figure 1.**
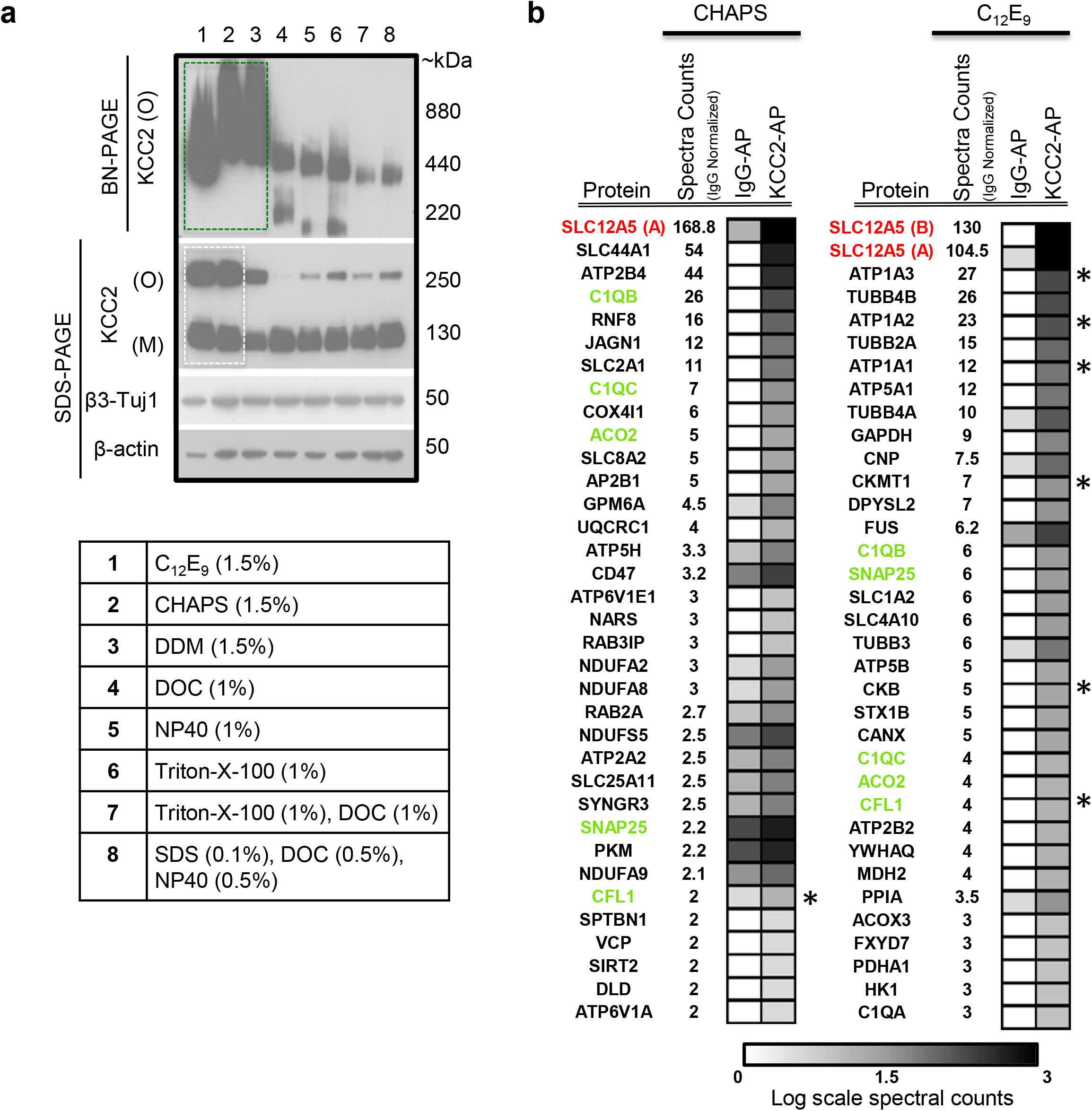
KCC2 multi-protein complexes can be extracted using native detergents. (a) BN-PAGE and SDS-PAGE separation of solubilized membrane fractions prepared from ~P50 mouse brain, using the detergents listed in the associated table. Protein separations were Western-blotted and probed with antibodies indicated on the left. O, oligomer; M, monomer. Blots are representative of two independent biological replicates. (b) Comparison of the top 35 proteins identified with high confidence in C-terminal KCC2 antibody immunoprecipitations from CHAPS-based or C_12_E_9_-based membrane extractions. IgG-AP immunoprecipitations were performed as a control. Heat maps represent log scale spectral counts of individual proteins per condition, expressed relative to global spectral counts. Unique peptides corresponding to KCC2 (indicated in red font) were most abundant in both conditions, confirming the specificity of the C-terminal antibody. Previously identified KCC2 interacting partners are identified by asterisks. Proteins in green represent those that commonly co-precipitated with KCC2 regardless of the detergent extraction. The following source data and figure supplements are available for Figure1: Figure 1 – Figure Supplement 1: Workflow to enrich KCC2 complexes Figure 1 – Figure Supplement 2:. SDS-PAGE separation of solubilized membrane fractions Figure 1 – Source Data 1: Proteins enriched in KCC2-AP using CHAPS vs C12E9

To determine which of these two detergents was optimal for our subsequent full-scale proteomic analysis, we performed AP-MS to compare the efficacy of C_12_E_9_ versus CHAPS-solubilized membrane fractions. Immunopurification was performed on membrane fractions prepared from adult (P50) wild-type (WT) mouse brain, using a well-validated commercially available C-terminal KCC2 antibody (Williams et al., 1999; Gulyas et al., 2001; Woo et al., 2002; Mahadevan et al., 2014) and a control IgG antibody. In both detergent conditions, we recovered maximum peptides corresponding to KCC2 from the KCC2 pull downs (KCC2-AP), in comparison to the control IgG pull downs (IgG-AP), confirming the specificity of the C-terminal KCC2 antibody (Figure 1b **and** Figure 1 – **Source Data 1**). However, upon further examination, two key pieces of evidence indicated that C_12_E_9_-based conditions are optimal for proteomic analysis of native KCC2. First, we observed peptides corresponding to both KCC2 isoforms-a and -b; in C_12_E_9_-based samples, but we could only detect peptides corresponding to KCC2 isoform-a in KCC2-APs from CHAPS-based samples. Second, we observed a higher enrichment of peptides corresponding to previously identified KCC2 interactors belonging to the family of Na^+^/K^+^ ATPases (ATP1A1-3), and the family of creatine kinases (CKB, CKMT1), and CFL1 in the KCC2-AP from C_12_E_9_-based samples. Based on these results we concluded that C_12_E_9_-based solubilization conditions yield more KCC2-specific binding partners and fewer IgG-specific binding partners compared to CHAPS, and thus provide a higher stringency for KCC2 AP-MS. Thus we performed all subsequent proteomic analysis of native KCC2 on samples solubilized with C_12_E_9_.

### Multi-epitope (ME) proteomic analysis of KCC2 complexes in the developing and mature brain

KCC2b is the most abundant isoform in the mature brain(Uvarov et al., 2009; Markkanen et al., 2014), and the isoform largely responsible for the extrusion of intracellular Cl^−^, and thus the shift from excitatory to inhibitory GABA during early postnatal development (Kaila et al., 2014). To focus our proteomic analysis on KCC2b we used a multi-epitope approach that allowed us to distinguish the KCC2 isoforms (Figure 2a). The C-terminal antibody recognizes both isoforms (Uvarov et al., 2007, 2009; Markkanen et al., 2014), so we chose to use another antibody that is specifically raised against the unique N-terminal tail of the KCC2b isoform. Lastly, we used a phosphospecific antibody for serine 940 (pS940), as phosphorylation of this residue increases KCC2 surface expression and/or transporter function (Lee et al., 2007, 2011). We validated these three KCC2 antibodies (C-terminal, N-terminal, and pS940) for KCC2-immunoenrichment (Figure 2 – Figure Supplement 1). Moreover, by taking a multi-epitope approach we significantly increased the likelihood of detecting KCC2 interactions that may be missed during single-epitope AP approaches. We performed 25 rounds of AP-MS on both developing (P5) and mature/adult (P50) WT mouse brain (Figure 2 – **Source Data 1**). We could not use KCC2-knockout brains since these animals die at birth, so as an alternative we used a mock IP for each sample condition in the absence of the KCC2 antibody using parallel preimmunization immunoglobulin (IgG/IgY) as negative controls. We obtained 440 potential KCC2 protein interactors with 99% confidence and a 1% false discovery rate. We identified KCC2 peptides spanning the entire sequence of KCC2 with ~44% sequence coverage, primarily at the C- and N-terminal tails (Figure 2b); and in both the developing and mature brain, KCC2 peptides were the most abundant peptides identified in the KCC2-IPs (Figure 2c). While the KCC2 C-terminal antibody recovered peptides from both isoforms of KCC2, the N-terminal KCC2b-specific antibody did not recover any KCC2a isoform-specific peptides, indicating the specificity of the antibodies used, and the success of KCC2-immunoenrichment in our AP-assays.

**Figure 2.**
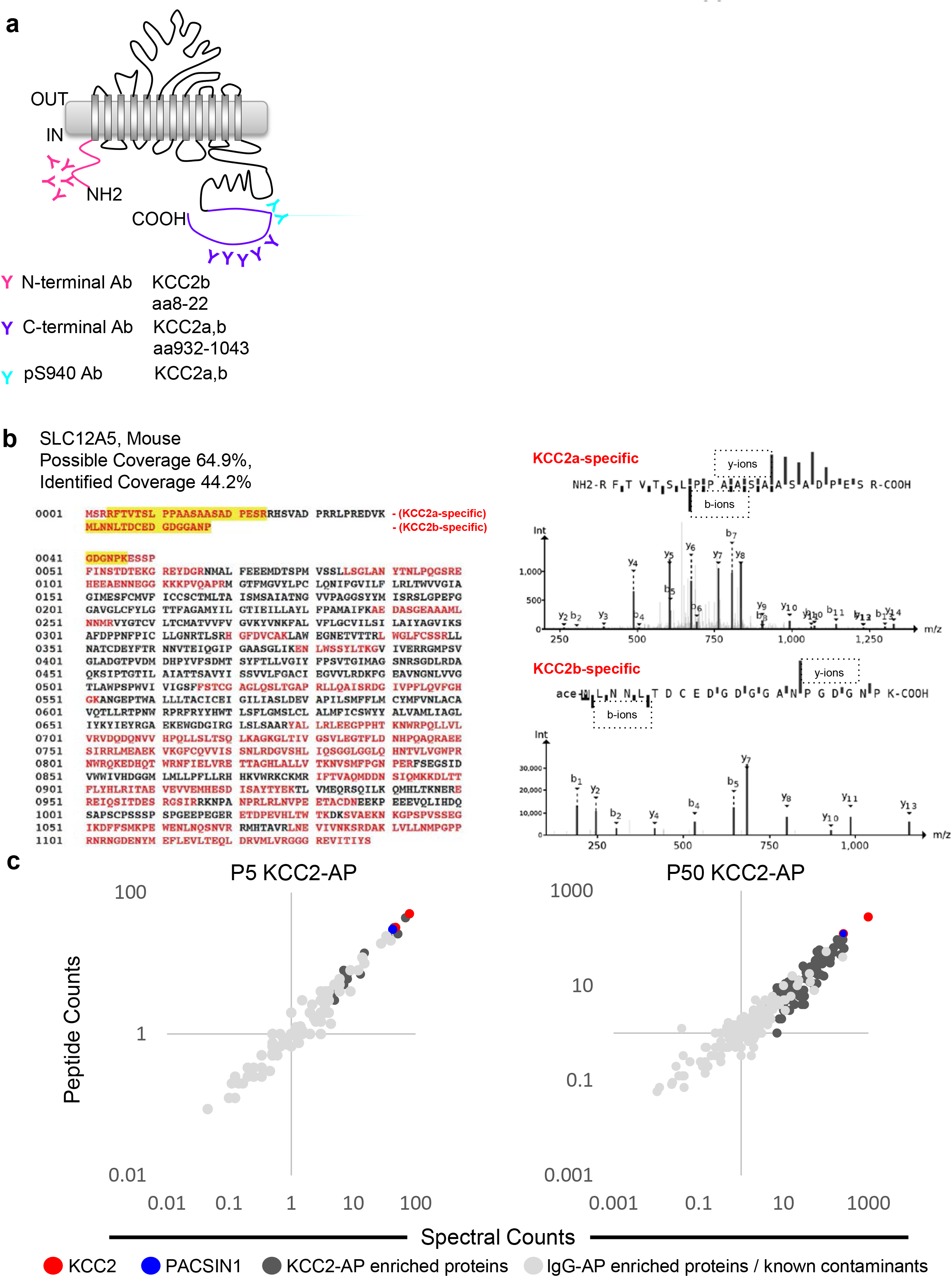
Multi-epitope AP identifies native-KCC2 protein constituents in mouse brain. **(a)** Schematic of the locations of anti-KCC2 antibodies. **(b)** The primary KCC2 amino acid sequence. Red indicates the protein coverage of KCC2 identified by MS analysis; yellow indicates unique coverage for KCC2a and KCC2b isoforms. MS/MS-spectra of peptides unique for KCC2a and KCC2b. Right: the MS/MS ion fragmentation of the corresponding amino acid sequence is indicated above the spectra. **(c)** Spectral and peptide count plots of proteins in AP with all three anti-KCC2 antibodies in developing brain membrane fractions (P5, left) and adult brain membrane fractions (P50, right). Peptide and spectral counts are normalized (anti-KCC2/IgG) and plotted on a log scale. Red circles - highly enriched KCC2 bait. Blue circles - highly enriched PACSIN1 target peptides. Dark-grey circles - top proteins enriched with KCC2-AP in comparison to IgG control-AP. Light-grey circles - proteins enriched in IgG control-AP in comparison to KCC2-AP and known spurious interactors. The following source data and figure supplements are available for Figure 2: Figure 2 – Figure Supplement 1. Validation of KCC2 antibodies used for immunodepletion. Figure 2 – Source Data 1: MS Replicates.

### The KCC2 interactome

To build the KCC2 interactome, all potential KCC2 protein interactors were filtered according to their spectral count enrichment in the KCC2-APs, and normalized to IgG IPs. In the first pass filter grouping, we included proteins with at least 2 unique peptides and peptide-spectrum matches and a 3-fold increase in KCC2 spectral counts in the KCC2-AP in comparison to IgG-AP (Figure 3 **– Source Data 1**). This yielded ~75 high-confidence, putative-KCC2 partners. In the second pass filter grouping, we identified additional high-confidence putative-KCC2 partners by including those with only 1 unique peptide, or less than 3-fold KCC2-AP enrichment, if they met one of the following criteria: (a) the protein was a previously validated KCC2 physical/functional interactor; (b) the protein family already appeared in the first-pass filter; (c) the protein appeared as a single-peptide interactor across multiple experiments (e.g. multiple antibodies, or in lysates from both age timepoints). Including these additional proteins from the second pass filtering yielded 186 putative-KCC2 partners. We next eliminated the 36 proteins that have been previously identified as commonly occurring spurious interactors in LC/MS experiments as indicated in the CRAPome database (Figure 3 **– Source Data 2**). (Mellacheruvu et al., 2013). Lastly, we added 31 proteins that have been previously established as KCC2-physical/functional partners but were not identified in our present LC-MS assay (Figure 3 **– Source Data 3**). By applying these filtering criteria and processes, we established a total list of 181 proteins in the KCC2 interactome (Figure 3 – Figure Supplement 1). More than half of these KCC2 interactors were exclusively enriched in KCC2-APs from the mature brain (85 proteins, ~57% overlap), while approximately one-third (41 proteins, ~27% overlap) were identified across both the developing and mature brain (Figure 3 – Figure Supplement 2). Only relatively small percentages were exclusively enriched in the developing brain (24 proteins, ~16% overlap).

**Figure 3.**
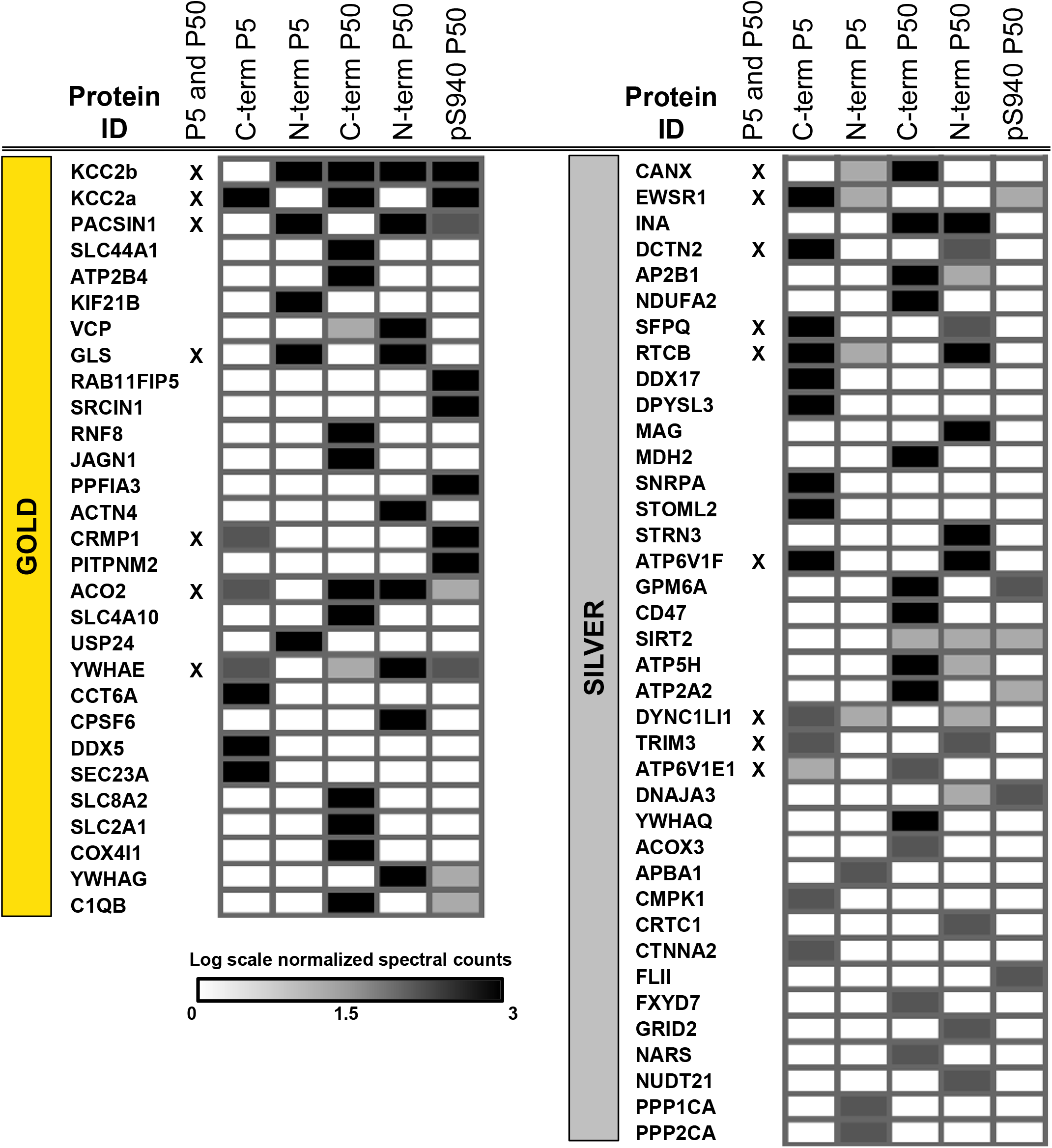
ME-AP reveals distinct KCC2 constituents in developing and mature brain. Summary of the top 70 proteins identified with high confidence across KCC2-ME AP in the developing and mature brain (3-fold spectral enrichment in KCC2-AP in comparison with IgG-AP). Heat map represents log scale spectral counts of individual proteins per antibody condition, expressed relative to global spectral counts. See Table 1 for a list of the transmembrane and soluble KCC2 interactors. The following source data and figure supplements are available for Figure 3: Figure 3 – Figure Supplement 1: The SLC12A5/KCC2 interactome Figure 3 – Figure Supplement 2: ME-AP proteomics identify the protein constituents of native KCC2 Figure 3 – Figure Supplement 3: Workflow for curating the KCC2 interactome Figure 3 – Source Data 1: Spectra-peptide counts for KCC2-AP and IgG-AP. Figure 3 – Source Data 2: CRAPome spurious interactors. Figure 3 – Source Data 3: Previously ID KCC2 interactors. Figure 3 – Source Data 4: KCC2 interactors identified in AP/MS categorized into gold, silver and bronze

We segregated the 181 protein KCC2 interactome into high-confidence (gold), moderate-confidence (silver), or lower confidence (bronze) putative KCC2-interactors (Figure 3, Table 1 **and** Figure 3 **– Source Data 4**). This tri-category segregation was based on the largest probability of a bait-prey pair across all replicate purifications, as indicated by the MaxP score (Choi et al., 2012). Gold KCC2-partners were those with normalized spectral count enrichments ≥5 and a - MaxP SAINT score ≥0.89. Silver KCC2-partners were those with normalized spectral count enrichments between 3 and 5, and a MaxP score between 0.7 and 0.89. Bronze KCC2-parterns were all remaining proteins that were not designated as Gold or Silver. Gold, Silver, and Bronze proteins were all included in subsequent network analysis.

**Table 1.**
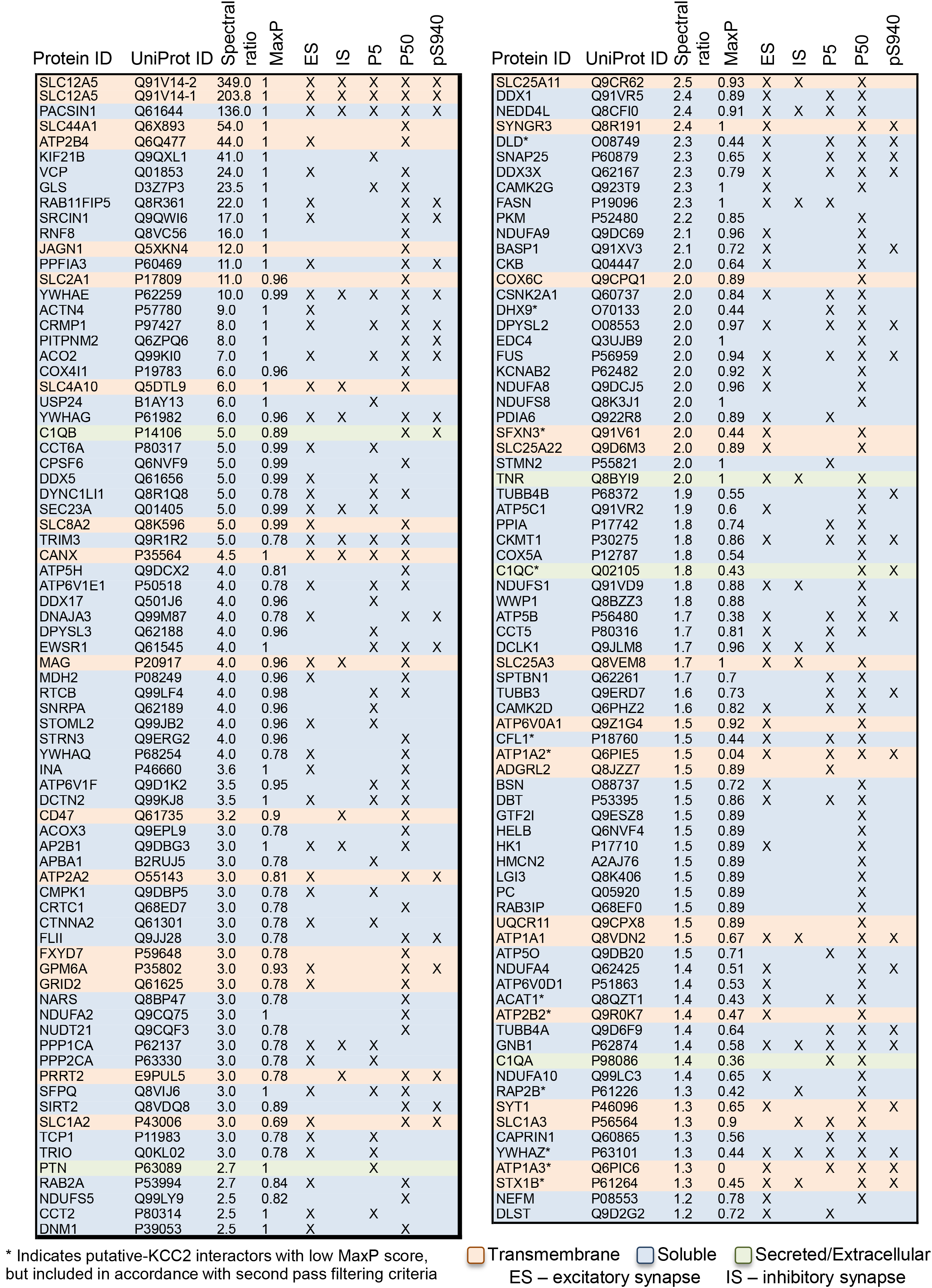

### Members of KCC2 interactome are highly represented at excitatory synapses

To interpret the potential functional role of KCC2-protein interactors we first segregated them based on their abundance at excitatory and inhibitory synapses. To perform this analysis we mapped the KCC2 interactome to the excitatory synapse-enriched postsynaptic density (PSD) proteome (Collins et al., 2006), or the inhibitory synapse-enriched proteomes (iPSD, GABA_A_R, GABA_B_R, NLGN2, and GlyR) (Heller et al., 2012; Del Pino et al., 2014; Kang et al., 2014; Nakamura et al., 2016; Schwenk et al., 2016; Uezu et al., 2016). Interactome mapping revealed that ~43% of proteins in the KCC2 interactome (77/181) were exclusively enriched at excitatory synapses, while only ~2% of proteins (4/181) were exclusively enriched at inhibitory synapses (Figure 4 a,b). However, ~15% proteins (28/181) were mapped to both excitatory and inhibitory synapses, while ~39% proteins (71/181) did not map to either synapse.

**Figure 4.**
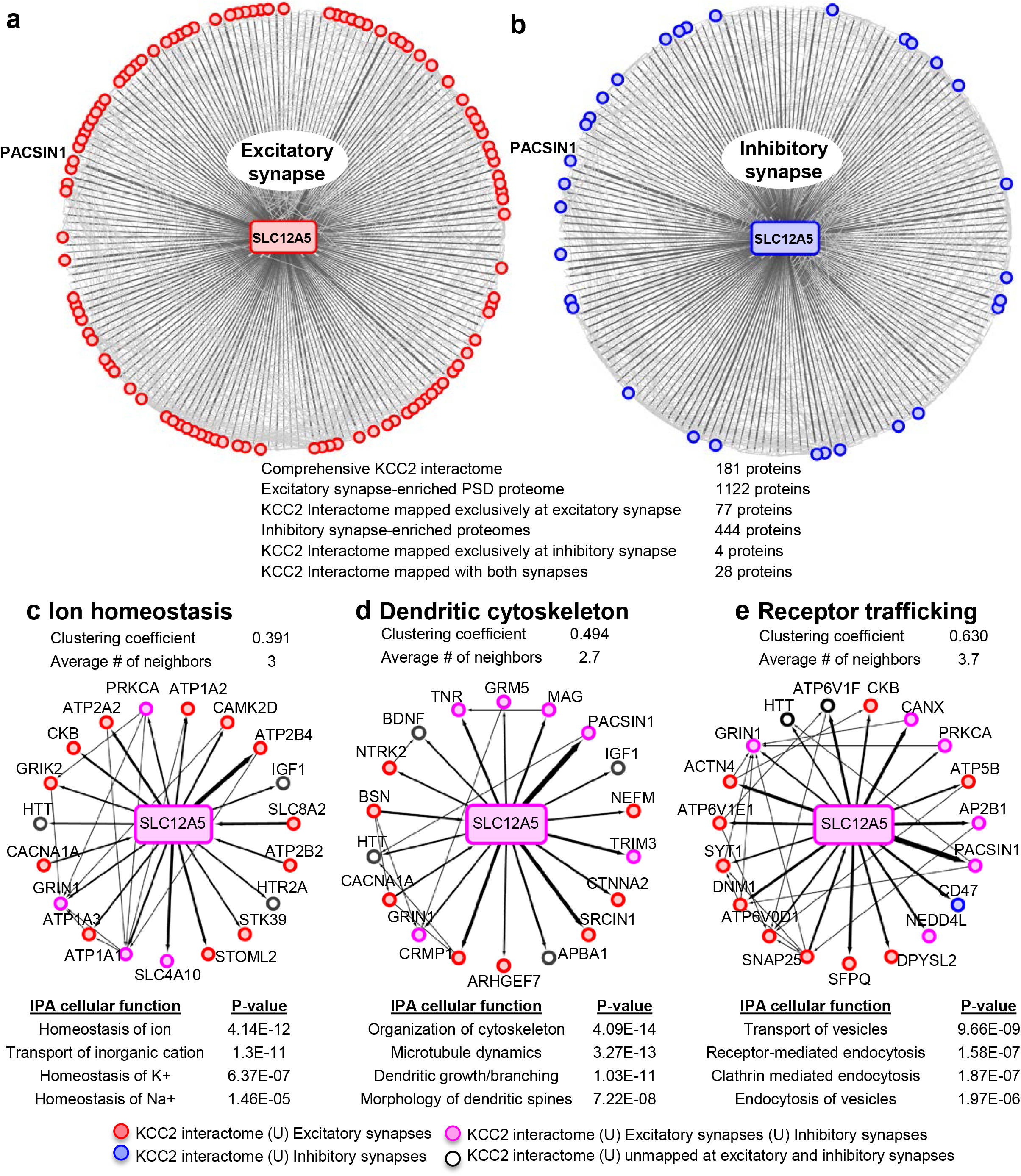
Members of the KCC2 interactome are highly represented at excitatory synapses. **(a)** The KCC2 interactome mapped to the excitatory synapse-enriched postsynaptic density proteome. Pink circles indicate proteins mapped to excitatory synapses, thickness of the edge denotes the number of spectral enrichment (KCC2/IgG) from the log scale. **(b)** Similar to a, but for the KCC2 interactome mapped to the inhibitory synapse-enriched proteomes for GABA_A_Rs, GABA_B_Rs, NLGN2, and GlyRs. **(c)** IPA revealing members of the KCC2 ME-AP involved in ion homeostasis; the thickness of the line represents the spectral enrichment (KCC2/IgG). **(d)** Similar to c, but for proteins involved in dendritic cytoskeleton rearrangement. **(e)** Similar to c, but for proteins involved in receptor recycling/endocytosis/trafficking. Source data for Figure 4 include: Figure 4 – Source Data 1: Ingenuity Pathway Anaysis.

To further examine the KCC2 interactome based on cellular functions we performed an Ingenuity Pathway Analysis (IPA) to segregate the KCC2-interactors into highly enriched Gene Ontology (GO) classes. Performing this IPA analysis revealed that KCC2 partners segregate into multiple cellular and molecular functional nodes, which we then combined into three broad categories that collectively had high p values: ion homeostasis, dendritic cytoskeleton rearrangement, and receptor trafficking (Figure 4 c–e; Figure 4 **– Source Data 1**). KCC2 has been previously associated with both ion homeostasis and dendritic spine morphology, and consistent with this previous work we identified previously characterized KCC2 functional or physical interactors, including subunits of the sodium/potassium (Na^+^/K^+^) ATPase, including the previously characterized KCC2 interactor ATP1A2 (Ikeda et al., 2004), and Cofilin1, which was recently demonstrated to be important for KCC2-mediated plasticity at excitatory synapses (Chevy et al., 2015; Llano et al., 2015). The third category, receptor trafficking, has a denser network (clustering coefficient of 0.63 and an average of ~3.7 neighbors) in comparison to the other networks, suggesting a tight link between KCC2 and proteins in this node. Notably, this analysis revealed multiple novel putative-KCC2 partners, including PACSIN1, SNAP25, RAB11FIP5, CK2, Dynm1 and AP2. All of these novel putative interactors have established functions in membrane recycling and/or trafficking of glutamate receptor subunits (Carroll et al., 1999; Lee et al., 2002; Vandenberghe et al., 2005; Pérez-Otaño et al., 2006; Selak et al., 2009; Sanz-Clemente et al., 2010; Anggono et al., 2013; Bacaj et al., 2015). In order to determine the spatiotemporal expression profiles of the KCC2 interactome, we first performed transcriptomic analysis and hierarchical clustering of high-resolution human brain RNAseq data (available at (http://brain-map.org). We observed that *SLC12A5* mRNA is expressed with several members members of the receptor trafficking node in the hippocampus (Figure 5a), amygdala, striatum, thalamus, cerebellum and cortex (Figure 5 – Figure Supplement 1; Figure 5 **– Source Data 1**).

**Figure 5.**
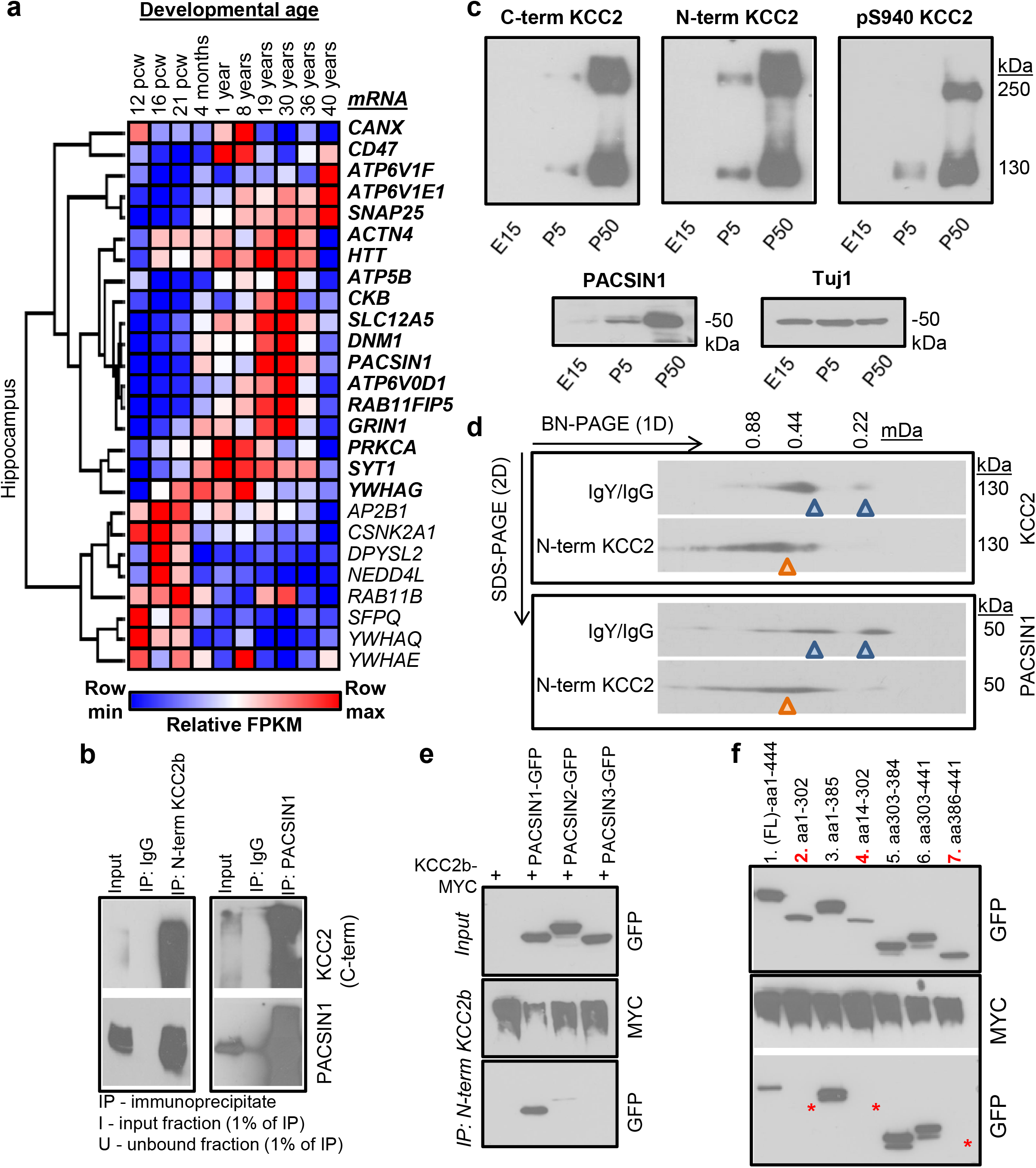
Characterization of the PACSIN1-KCC2 interaction. **(a)** Spatiotemporal expression patterns of SLC12A5 and members of the receptor trafficking node in the human brain **(b)** Native KCC2 complexes from C_12_E_9_-solubilized whole-brain membrane fractions immunoprecipitated with IgY or anti-N-term KCC2 (left) and IgG or anti-PACSIN1 (right), and immunoblotted with C-term KCC2 and PACSIN 1 antibodies. **(c)** Western blot of KCC2 and PACSIN1 during development from C_12_E_9_-solubilized hippocampal membrane fractions (probed with the antibodies indicated above). **(d)** Antibody-shift assay followed by 2D-BN-PAGE separation using C_12_E_9_-solubilized whole-brain membrane fractions, incubated with antibodies indicated on left. **(e)** Coimmunoprecipitation experiments performed in COS7 cells transfected with myc-tagged KCC2b and GFP-tagged PACSIN1/2/3 constructs, immunoprecipitated with anti-N-term KCC2. **(f)** Immunoblot of immunoprecipitates from transfected COS7 cell lysates. * indicate the lanes where PACSIN1 lacks the variable region between ~aa325-383. independent biological replicates: 5e = 4; 5f = 3; 5b,c,d = 2. The following source data and figure supplements are available for Figure 5: Figure 5 - Source Data 1: Allan Brain Atlas data. Figure 5 – Supplement Figure 1: Spatiotemporal expression patterns of *SLC12A5* and members of receptor trafficking node. Figure 5 – Supplement Figure 2: The primary amino acid sequence coverage of PACSIN1

In order to independently validate the KCC2 interactome, we proceeded to biochemical and functional analysis. We focused this validation analysis on proteins in the receptor trafficking category for two reasons: (i) the most abundant putative-KCC2 partner, PACSIN1 (PKC and CK2 substrate in neurons; also called as Syndapin1) is present in the receptor trafficking node; and (ii) the tightest KCC2-subnetwork exists in receptor trafficking node, indicating a dense interconnectivity between these proteins.

### PACSIN1 is a novel native-KCC2 binding partner

To biochemically and functionally validate our KCC2 interactome we chose to focus on the putative KCC2-PACSIN1 interaction. The rationale for this selection was based on the following: (1) PACSIN 1 is the most abundant KCC2 interactor in our analysis, with a high normalized spectral count ratio and a high MaxP score, and with extensive amino acid sequence coverage (**Figure 5 – Figure Supplement 2**); (2) PASCIN1 is a substrate for PKC, and PKC is a key regulator of KCC2 (Lee et al., 2007); (3) PASCIN1 is a substrate for CK2, and our analysis revealed CK2 as a putative KCC2-interactor; (4) PACSIN1 is abundant at both excitatory and inhibitory synapses; and (5) PACSIN1 was identified as an abundant KCC2 interactor using multiple antibodies (N-terminal and pS940).

To independently verify whether KCC2 associated with PACSIN1, we performed a co-immunoprecipitation assay from adult whole-brain native membrane preparations. We found that anti-KCC2b antibodies, but not control IgY antibodies, co-immunoprecipitated with PACSIN1 (**Figure 5b**). Using a previously well-validated PACSIN1 antibody (Anggono et al., 2013) we confirmed this interaction in the reverse direction, indicating the existence of a KCC2-complex with PACSIN1 *in* vivo (**Figure 5b**). Consistent with our ability to co-immunoprecipitate KCC2 and PACSIN1, we found that the expression profiles of KCC2 and PACSIN1 are temporally aligned in the mouse brain (**Figure 5c**). To determine whether native-KCC2 complexes are stably associated with PACSIN1, we performed an antibody-shift assay coupled with two-dimensional BN-PAGE (2D BN-PAGE), which is a strategy that has been used previously to examine the native assemblies of several transmembrane protein multimeric complexes (Schwenk et al., 2010, 2012), including that of native-KCC2 (Mahadevan et al., 2014). Using this approach, we first verified that the addition of N-terminal KCC2b antibodies could shift a proportion of native-KCC2 to higher molecular weights, in comparison to IgY control antibodies (**Figure 5d**). Next, we observed that this antibody-induced shift in native-KCC2b using N-terminal antibody also shifted a population of native-PACSIN1 to comparable higher molecular weights (**Figure 5d**). Collectively, these experiments establish native-PACSIN1 as a novel KCC2-binding partner in whole brain tissue.

The PACSIN family of proteins contains 3 members that share ~90% amino acid identity (Modregger et al., 2000). PACSIN1 is neuron-specific and is broadly expressed across multiple brain regions; PACSIN2 is ubiquitous and is abundant in cerebellar Purkinje neurons (Anggono et al., 2013; Cembrowski et al., 2016), and PACSIN3 is restricted to muscle and heart (Modregger et al., 2000). To determine which members of the PACSIN family binds to KCC2, we transfected PACSIN constructs (Anggono et al., 2013), with myc-KCC2b in COS-7 cells and performed co-immunoprecipitation. We observed that KCC2 robustly associates with PACSIN1, weakly interacts with PACSIN2, and does not interact with PACSIN3 (**Figure 5e**). PASCIN1 contains a membrane-binding F-BAR domain, a SH3 domain that binds to phosphorylated targets, and a VAR (variable) region (Kessels and Qualmann, 2004, 2015). In order to determine the PACSIN1 region that is required for KCC2 binding we repeated our co-immunoprecipitation assays in COS-7 cells, but this time we used previously characterized PACSIN1 deletion constructs (Anggono et al., 2013) (**Figure 5f**). We discovered that removing either the SH3 or the F-BAR region did not disrupt the KCC2:PACSIN1 interaction, indicating that they are not necessary for KCC2 binding. In an analogous result, neither the SH3 domain nor the F-BAR domain could interact with KCC2. However, KCC2 robustly co-precipitated with PACSIN1 when the VAR region was co-expressed with KCC2, indicating that the VAR region is sufficient to mediate the KCC2 interaction

### PACSIN1 is a negative regulator of KCC2 expression and function in hippocampal neurons

KCC2 dysregulation has emerged as a key mechanism underlying several brain disorders including seizures (Fiumelli et al., 2013; Stödberg et al., 2015; Saitsu et al., 2016), neuropathic pain (Coull et al., 2003), schizophrenia (Tao et al., 2012), and autism spectrum disorders (ASD) (Cellot and Cherubini, 2014; Tang et al., 2015). However there are currently no existing KCC2 enhancers approved for clinical use, and thus there is a critical need to identify novel targets for the development of KCC2 enhancers. To determine whether PACSIN1 may be a potential target for regulating KCC2 function, we assayed for KCC2 function following PACSIN1 knockdown. We chose to assay for the canonical KCC2 function of Cl^−^ extrusion, as the loss of Cl^−^ homeostasis and thus synaptic inhibition, is causal for several neurological disorders (Coull et al., 2003; Huberfeld et al., 2007; Tao et al., 2012; Cellot and Cherubini, 2014; Toda et al., 2014; Kahle et al., 2014; Puskarjov et al., 2014; Stödberg et al., 2015; Banerjee et al., 2016; Saitsu et al., 2016; Tang et al., 2016). We assayed KCC2-mediated Cl^−^ extrusion by performing whole cell recordings of the reversal potential for GABA (E_GABA_) in cultured hippocampal neurons while loading the neuron with Cl^−^ to drive KCC2 transport. When neurons were virally transfected with a previously validated PASCIN1 silencing shRNA construct (Anggono et al., 2013), E_GABA_ was hyperpolarized relative to neurons transfected with the control shRNA construct (**Figure 6a,b**; control shRNA: −28.62 ± 3.07 mV, n = 9; PACSIN1 shRNA: −37.86 ± 1.73 mV, n = 11; t(18)=2.744, p = 0.013), with no significant change in the GABA_A_R conductance (**Figure 6a,c**; control shRNA: 6.93 ± 1.32 mV, n = 9; PACSIN1 shRNA: 12.96 ± 2.71 mV, n = 11; t(18)=1.86, p = 0.079). In addition, we performed gramicidin-perforated patch clamp recordings to maintain Cl^−^ gradients and a significant hyperpolarizing shift in E_GABA_ compared to whole cell recordings (**Figure 6d**; whole-cell recordings n = 11; gramicidin recordings n = 6; t(15)=4.021, p=0.001). Thus, PACSIN1 silencing increases KCC2-mediated Cl^−^ extrusion in neurons, which we predicted might be due to an increase in KCC2 expression. To test our prediction we performed immunofluorescent staining of endogenous KCC2 in cultured hippocampal neurons transfected with either shRNA-control or shRNA-PACSIN1. We observed a significant increase in KCC2 fluorescence in neurons expressing PASCIN1-shRNA in comparison with control shRNA (**Figure 6e**; control shRNA: 58.86 ± 2.53 A.U., n = 32; PACSIN1 shRNA: 74.05 ± 2.53 A.U., n = 32; t(31)=5.272, p < 0.0001). Taken together, our electrophysiological recordings and immunofluorescence results demonstrate that a reduction in PACSIN1 results in increased KCC2 expression and an increase in the strength of inhibition (hyperpolarization of E_GABA_). If PACSIN1 is a *bona fide* negative regulator of KCC2 expression, then overexpressing PACSIN1 should produce a reduction in KCC2 expression. To test this prediction we performed immunofluorescent staining of endogenous KCC2 in cultured hippocampal neurons transfected with either eGFP or PASCIN1-eGFP. We observed a remarkable loss of KCC2 immunofluorescence when PACSIN1 was overexpressed in comparison to control eGFP (**Figure 6f**; control eGFP: 62.1 ± 2.7 A.U., n = 23; PACSIN1-eGFP: 11.31 ± 3.17 A.U., n = 16; t(37)=12.13, p < 0.0001).

**Figure 6.**
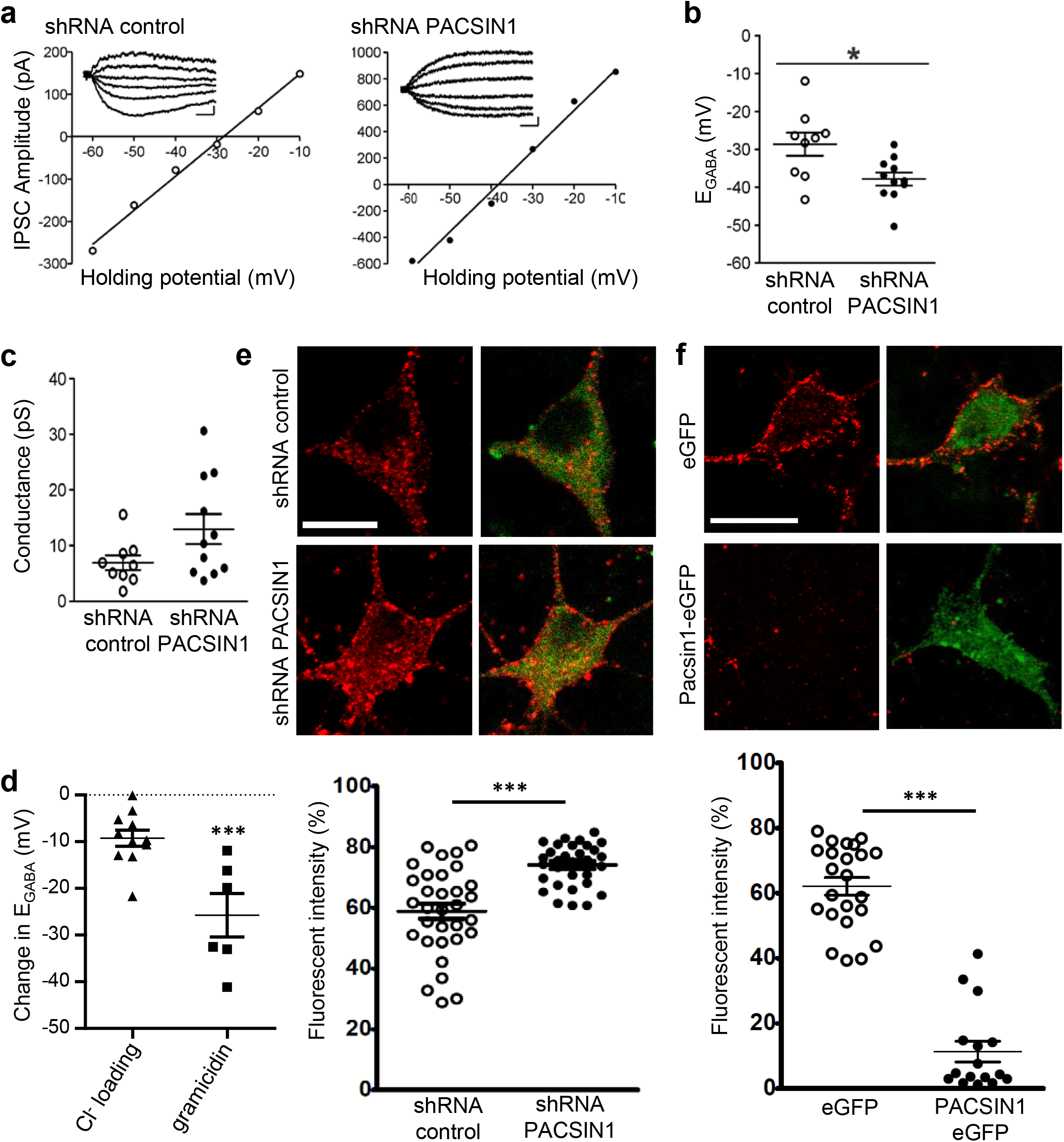
PACSIN1 is a negative regulator of KCC2 expression and function. **(a)** Example IV curves measuring E_GABA_ using Cl^−^-loading through whole-cell configuration from cultured hippocampal neurons transduced with control shRNA (n=9) or PACSIN1 shRNA (n=11). Summary of **(b)** E_GABA_ and **(c)** synaptic conductance from all experiments similar to a (mean ± sem). **(d)** Change in E_GABA_ between whole-cell (n=11) and gramicidin-perforated (n=6) recordings (mean ± sem). **(e)** Example confocal microscopic immunofluorescent images from cultured hippocampal neurons transduced with control shRNA (n=32) or PACSIN1 shRNA (n=32) and stained with anti-KCC2 (red; scale bar, 10μm); green immunostain reports transfection. Below: summary of fluorescence intensities (mean ± sem). **(f)** Similar to e, except neurons were transduced with either control eGFP (n=23) or PACSIN1-eGFP (n=16). n values for all experiments on cultured neurons were obtained from a minimum of 3 independent sets of cultures. Statistical significance was determined using student’s t-tests (two-tailed); * p < 0.05, *** p < 0.001

## DISCUSSION

We determined that the mouse brain KCC2 functional interactome is comprised of 181 proteins; and by mapping the KCC2 interactome to excitatory and inhibitory synapse proteomes and performing ingenuity pathway analysis, we determined that KCC2 partners are highly enriched at excitatory synapses and form a dense network with proteins involved in receptor trafficking. We validated the KCC2-interactome by biochemically characterizing the interaction between KCC2 and the most abundant protein in the interactome, PACSIN1. Functional validation of the KCC2-PACSIN1 interaction revealed that PACSIN1 robustly and negatively regulates KCC2 expression. While ion channels and GPCRs are known to predominantly exist in large multi-protein complexes (Husi et al., 2000; Berkefeld et al., 2006; Collins et al., 2006; Müller et al., 2010; Schwenk et al., 2010, 2012, 2014, 2016; Nakamura et al., 2016; Pin and Bettler, 2016), similar studies on solute carrier proteins (transporters) are still in their infancy (Snijder et al., 2015). Based on the critical importance of SLC transporters as therapeutic targets in both rare and common diseases (César-Razquin et al., 2015; Lin et al., 2015), including that of KCC2 in human neurological diseases (Blaesse et al., 2009; Medina et al., 2014), our present study also fills a general gap in the field of CNS transporter proteomics.

We report that native-detergents C_12_E_9_ and CHAPS extract KCC2 isoforms differentially (**Figure 1b**). A common caveat of isoform-counting in shot-gun proteomic experiments such as ours, is the problem of protein inference (Nesvizhskii and Aebersold, 2005). While we were able to discriminate KCC2 isoforms-a and -b, based on the presence of their unique peptides in their N-terminus (**Figure 1c**), it is not possible to categorize the remaining peptides to either isoform a or b due to their extensive shared homology. Therefore targeted proteomics such as selected-reaction monitoring would be required to accurately establish the abundances of KCC2 isoforms. We also report that native-detergents C_12_E_9_ and CHAPS pull-down different subsets of proteins along with some common interactors (**Figure 1b**). It is intriguing to note that there were several putative KCC2-partners uniquely identified with this detergent (SLC44A1, ATP2B4, RNF8, JAGN1, CD47, SLC2A1 (**Figure 1b**). Although we did not perform exhaustive proteomics with CHAPS-based KCC2 extractions, because of the presence of these high-confidence proteins in the CHAPS-based KCC2 LC/MS, we did include these proteins in the KCC2-interactome. This demonstrates that detergent stabilities of KCC2 protein complexes are distinct, in line with other recent ion channel proteomic studies (Müller et al., 2010; Schulte et al., 2011; Schwenk et al., 2012).

Our KCC2 LC/MS identified previously established KCC2 proteins interactors, including ATP1A2 (Ikeda et al., 2004), CFL1 (Chevy et al., 2015; Llano et al., 2015) and CKB (Inoue et al., 2004, 2006), which add confidence to the validity of this interactome. We were initially surprised at the absence of other previously identified KCC2 interactors, including: Neto2 (Ivakine et al., 2013; Mahadevan et al., 2015), GluK2 (Mahadevan et al., 2014; Pressey et al., 2017), 4.1N (Li et al., 2007), beta-pix (Chevy et al., 2015; Llano et al., 2015), RCC1 (Garbarini and Delpire, 2008), or signaling molecules PKC (Lee et al., 2007), WNK, SPAK and OSR (Friedel et al., 2015). The absence of these previously identified interactors may be due to any of the following caveats, which have been well recognized in previous ion channel and GPCR proteomic studies: (1) these interactions may be weak, transient, mediated by posttranslational modifications (Schulte et al., 2011), or mediated by intermediary partners; (2) these interactions are under-represented because they are restricted to specific brain regions; (3) antibody-epitope binding knocked-off endogenous interactions; (4) despite using the C_12_E_9_-based solubilization strategy that is known to stabilize ion pumps and transporters (Romero, 2009; Babu et al., 2010; Ramachandran et al., 2013) particular interactions may be better preserved by other detergent conditions.

Single particle tracking of surface KCC2 has revealed that ~66% of KCC2 is located synaptically (Chamma et al., 2012, 2013). While the density of surface KCC2 was not reportedly different between excitatory and inhibitory synapses, KCC2 was shown to dwell longer at excitatory synapses. Our observation that KCC2 interacting proteins are primarily enriched at excitatory synapses in comparison to inhibitory synapses is in line with this increased confinement of KCC2 at excitatory synapses. The presence of KCC2 at excitatory synapses raises some interesting questions: How does KCC2–mediated Cl^−^ extrusion regulate hyperpolarizing inhibition if it is preferentially localized near excitatory synapses? Why are the KCC2 partners exclusive at the inhibitory synapses less represented when compared with excitatory synapses? One potential answer to both of these questions is that because of the difficulty in identifying components of inhibitory synapses our knowledge of the proteins present at these structures is incomplete. Despite the fact that our network mapping incorporated 444 proteins known to be enriched at inhibitory synapses (Heller et al., 2012; Del Pino et al., 2014; Kang et al., 2014; Nakamura et al., 2016; Schwenk et al., 2016; Uezu et al., 2016), it is possible that we identified a smaller representation of inhibitory synapse-specific KCC2 partners in our present study. Another possibility is that KCC2 ‘moon-lights’ between inhibitory and excitatory synapses, as previously suggested (Blaesse and Schmidt, 2014). Our interactome supports this hypothesis as we identified 28 proteins that are enriched at both synapses. However, future studies are required to systematically examine whether the KCC2 complexes containing these 28 proteins enriched at both loci are similar or distinct. While the notion that excitatory and inhibitory synapses are distinct structures is widely accepted, emerging evidence from cortex suggests this may not be strictly true (Chiu et al., 2013; Higley, 2014). Under circumstances where excitatory and inhibitory synapses are in close physical proximity, the molecular complex involving KCC2 and these moonlighting proteins are ideally placed to execute cell-intrinsic E/I balance regulation, a hypothesis stemming from our present study that requires rigorous experimental testing.

Ever since the first discovery that KCC2 participates in the regulation of dendritic structures (Li et al., 2007), several studies have demonstrated 4.1N as a critical mediator of this non-canonical transporter-independent KCC2 function (Horn et al., 2010; Gauvain et al., 2011; Chamma et al., 2013; Fiumelli et al., 2013). Recently however, additional molecular players underlying this phenomenon, including COFL1, and ARHGEF7 (Beta-pix) have been identified to interact with KCC2 (Chevy et al., 2015; Llano et al., 2015). In the present study, we identify diverse high confidence (Gold) cytoskeletal organizers belonging to distinct protein families such as CRMP proteins, SRCIN1, VCP, KIF21B, previously unsuspected to mediate KCC2-dependant non-canonical function. However the precise relation between KCC2-dependant non-canonical functions and these putative partners in not currently known.

PACSIN1 is a well-established endocytic adapter protein that regulates the surface expression of distinct glutamate (Anggono et al., 2006, 2013; Pérez-Otaño et al., 2006; Widagdo et al., 2016) and glycine receptors (Del Pino et al., 2014). We reveal PACSIN1 as a novel negative regulator of KCC2 expression in central neurons. We previously reported that native-KCC2 assembles as a hetero-oligomer that migrates predominantly above ~400kDa (Mahadevan et al., 2014, 2015). Similar to KCC2, native-PACSIN1 also migrates above ~400kDa (Kessels and Qualmann, 2006). Here we report that while *SLC12A5* and *PACSIN1* mRNA transcripts increase in parallel in multiple brain regions throughout development, PACSIN1 overexpression remarkably decreases total KCC2 abundance. How does PACSIN1 exist in a stable complex with KCC2 when it negatively regulates KCC2 expression? Since KCC2 and PACSIN1 are both dynamically regulated by phosphorylation and PKC (Anggono et al., 2006; Lee et al., 2007; Clayton et al., 2009; Kahle et al., 2013), we predict that upon KCC2 phosphorylation, PACSIN1 uncouples from KCC2 rendering it incapable of negatively regulating KCC2. Numerous pathological situations are associated with decreased KCC2 phosphorylation at Ser940 (Wake et al., 2007; Lee et al., 2011; Sarkar et al., 2011; Toda et al., 2014; Ford et al., 2015; Mahadevan et al., 2015; Silayeva et al., 2015; Leonzino et al., 2016; Mahadevan and Woodin, 2016), resulting in decreased transporter expression and/or function. It will be important to determine whether any of these neurological deficits stem from PACSIN1-mediated decreases in KCC2. In the present study, we demonstrate that PACSIN1 shRNA increases KCC2 expression and strengthens inhibition, indicating that PACSIN1 is a target for intervention to upregulate KCC2 during pathological states.

## MATERIALS AND METHODS

### Animals and approvals

All experiments were performed in accordance with guidelines and approvals from the University of Toronto Animal Care Committee and the Canadian Council on Animal Care. Animals of both sexes from wild-type mice, C57/Bl6 strain (Charles River Laboratories) were used throughout. Animals were housed in the Faculty of Arts and Science Biosciences Facility (BSF) in a 12h light: 12h d cycle, with 2–5 animals/cage.

### Detergents

All biochemical preparations and centrifugations were performed at 4 °C as previously described (Ivakine et al., 2013; Mahadevan et al., 2014, 2015). Systematic analysis of detergent solubility, and migration of native-KCC2 from crude membrane fractions were performed according to the workflow described in **Figure 1 – Figure Supplement 1**. The following 8 detergents (or detergent combinations) were used to solubilize whole brain membranes: C_12_E_9_ (1.5%), CHAPS (1.5%), DDM, (1.5%), DOC (1%), NP40 (1%), Triton-X-100 (1%), Triton-X-100 (1%) + DOC (1%), SDS (0.1%) + DOC (1%) + NP40 0.5%).

### Purification of KCC2 and in vivo co-immunoprecipitation

Mice (~P5, P50) were sacrificed, and brains were removed and homogenized on ice in PBS using a glass-Teflon homogenizer, followed by brief low-speed centrifugation. Soft-pellets were re-suspended in ice-cold lysis buffer [Tris·HCl, 50 mM, pH 7.4; EDTA, 1 mM; protease and phosphatase inhibitor mixture (Roche)], homogenized, and centrifuged for 30 minutes at 25,000 × g. Membrane pellets were resuspended in solubilization buffer (4Xw/v) [Tris·HCl, 50 mM, pH 7.4; NaCl, 150 mM; EDTA, 0.05 mM; selected detergent(s), and protease and phosphatase inhibitor mixture(Roche)], solubilized for 3 hours on a rotating platform at 4 °C, and centrifuged for 1 hour at 25,000 × g. For KCC2 and control co-immunoprecipitations, 20-100 μl GammaBind IgG beads were incubated on a rotating platform with the following antibodies (5-100 μg antibody) for 4 hours at 4 °C in cold 1X PBS:

- mouse polyclonal C-term KCC2 antibody, Neuromab #N1/12, RRID AB_10697875;
- rabbit polyclonal C-term KCC2 antibody, Millipore #07-432, RRID AB_310611;
- mouse monoclonal pS940 KCC2, Phosphosolutions #p1551-940, RRID AB_2492213;
- chicken polyclonal N-term KCC2b antibody (Markkanen et al., 2014);
- IgG/IgY control antibodies

Following antibody binding, 20 mM DMP (dimethyl pimelimidate, ThermoFisher 21667) in cold 1X PBS was used to crosslink antibodies with the beads, according to manufacturer’s instructions. The crosslinking reaction was stopped by adding 50 mM Tris·HCl to quench excess DMP, and the antibody-conjugated beads were thoroughly washed with the IP buffer. 1-10 mg of pre-cleared mouse brain membrane fractions were incubated with KCC2 or control antibody-conjugated beads on a rotating platform for 4 hours at 4 °C. After co-immunoprecipitation, the appropriate unbound fraction was saved for comparison with an equal amount of lysate to calculate the IP-efficiency (**Figure 2 – Figure Supplement 1**). The beads were washed twice with IP buffer containing detergent, and twice with IP-buffer excluding the detergent. The last wash was performed in 50 mM ammonium bicarbonate. Co-immunoprecipitation experiments for validating KCC2 and PACSIN1 was performed similar to the above procedure, in the absence of DMP-crosslinking. In a subset of validation experiments, anti-PACSIN1 antibody (Synaptic Systems #196002, RRID AB_2161839), was used for reverse co-IP. The break-down of LC/MS replicates are as follows:

- Optimization of LC/MS (**Figure 1**) using CHAPS and C_12_E_9_-solubilized membrane fractions were performed each with parallel IgG. (6XCHAPS KCC2) + (6XCHAPS IgG) + (2XC12E9 KCC2) + (2XC12E9 IgG) = 16 AP/MS using 5 μg C-terminal pan-KCC2 antibody, and 1mg of P50 membranes.
- LC/MS (**Figure 2,3**) using C_12_E_9_-solibilized membrane fractions were performed (with parallel IgG/IgY) as follows: (1XP50, C-term KCC2) + (1XP50, pS940 KCC2) + (1XP50, IgG) + (1XP50, N-term KCC2) + (1XP50, IgY) + (1XP5, C-term KCC2) + (1XP5, IgG) + (1XP5, N-term KCC2) + (1XP5, IgY) = 9 AP/MS using 100 μg KCC2 antibody, and 10 mg of membrane.

### Mass spectrometry

Mass spectrometry for the creation of the KCC2 interactome (**Figure 2, 3**) was performed at the SPARC Biocentre at SickKids Research Institute (Toronto, Ontario). Mass spectrometry for the determination of optimal detergents for native KCC2 extraction (**Figure 1**) was performed in the lab of Dr. Tony Pawson at the Lunenfeld-Tanenbaum Research Institute (LTRI), Mount Sinai Hospital (Toronto, ON) and in the CBTC (University of Toronto). Specific details on the individual experiments performed in each facility is located in **Figure 2 – Source Data 1**.

For all MS experiments, proteins were eluted from beads by treatment with double the bead volume of 0.5 M ammonium hydroxide (pH 11.0), and bead removal by centrifugation; this procedure was repeated 2x. The combined supernatants were dried under vacuum, reduced with DTT, and the free cysteines were alkylated with iodoacetamide. The protein concentration was measured, and trypsin was added at a ratio of 1:50; digestion occurred overnight at 37 °C. The peptides were purified by C18 reverse phase chromatography on a ZipTip (Millipore). Specifics of the MS in the three facilities are below: <u>SPARC</u>: The peptides were analyzed on a linear ion trap-Orbitrap hybrid analyzer (LTQ-Orbitrap, ThermoFisher, San Jose, CA) outfitted with a nanospray source and EASY-nLC split-free nano-LC system (ThermoFisher, San Jose, CA). Lyophilized peptide mixtures were dissolved in 0.1% formic acid and loaded onto a 75 μm x 50 cm PepMax RSLC EASY-Spray column filled with 2 μM C18 beads (ThermoFisher San, Jose CA) at a pressure of 800 BAR. Peptides were eluted over 60 min at a rate of 250 nl/min using a 0 to 35% acetonitrile gradient in 0.1% formic acid. Peptides were introduced by nano electrospray into an LTQ-Orbitrap hybrid mass spectrometer (Thermo-Fisher). The instrument method consisted of one MS full scan (400–1500 m/z) in the Orbitrap mass analyzer, an automatic gain control target of 500,000 with a maximum ion injection of 200 ms, one microscan, and a resolution of 120,000. Ten data-dependent MS/MS scans were performed in the linear ion trap using the ten most intense ions at 35% normalized collision energy. The MS and MS/MS scans were obtained in parallel fashion. In MS/MS mode automatic gain control targets were 10,000 with a maximum ion injection time of 100 ms. A minimum ion intensity of 1000 was required to trigger an MS/MS spectrum. The dynamic exclusion was applied using a maximum exclusion list of 500 with one repeat count with a repeat duration of 15 s and exclusion duration of 45 s.

<u>CBTC</u>: Orbitrap analyzer (Q-Exactive, ThermoFisher, San Jose, CA) outfitted with a nanospray source and EASY-nLC nano-LC system (ThermoFisher, San Jose, CA). Lyophilized peptide mixtures were dissolved in 0.1% formic acid and loaded onto a 75μm x 50cm PepMax RSLC EASY-Spray column filled with 2μM C18 beads (ThermoFisher San, Jose CA) at a pressure of 800 Bar. Peptides were eluted over 60 min at a rate of 250nl/min using a 0 to 35% acetonitrile gradient in 0.1% formic acid. Peptides were introduced by nanoelectrospray into the Q-Exactive mass spectrometer (Thermo-Fisher). The instrument method consisted of one MS full scan (400–1500 m/z) in the Orbitrap mass analyzer with an automatic gain control target of 1e6, maximum ion injection time of 120 ms and a resolution of 70,000 followed by 10 data dependent MS/MS scans with a resolution of 17,500, an AGC target of 1e6, maximum ion time of 120ms, and one microscan. The intensity threshold to trigger a MS/MS scan was set to 1.7e4. Fragmentation occurred in the HCD trap with normalized collision energy set to 27. The dynamic exclusion was applied using a setting of 10 seconds.

<u>LTRI</u>: Nano-LCMS using a home-packed 0.75 μm × 10cm C18 emitter tip (Reprosil-Pur 120 C18-AQ, 3 μm). A Nano LC-Ultra HPLC system (Eksigent) was coupled to an LTQ Orbitrap Elite (ThermoFisher) and samples were analyzed in data-dependent acquisition mode. A 60000 resolution MS scan was followed by 10 CID MS/MS ion trap scans on multiply charged precursor ions with a dynamic exclusion of 20 seconds. The LC gradient was delivered at 200nl/minute and consisted of a ramp of 2-35% acetonitrile (0.1% formic acid) over 90minutes, 35-80% acetonitrile (0.1% formic acid) over 5 minutes, 80% acetonitrile (0.1% formic acid) for 5 minutes, and then 2% acetonitrile for 20 minutes.

### Analysis of mass spectra and protein identification

All MS/MS samples were analyzed using Sequest (Thermo Fisher Scientific, San Jose, CA, USA; version 1.4.0.288) and X! Tandem (The GPM, thegpm.org; version CYCLONE (2010.12.01.1)). Sequest was set up to search Uniprot-mus+musculus_reviewed_Oct172015.fasta (unknown version, 25231 entries) assuming the digestion enzyme trypsin. X! Tandem was set up to search the Uniprot-mus+musculus_reviewed_Oct172015 database (unknown version, 25248 entries) also assuming trypsin. Sequest and X! Tandem were searched with a fragment ion mass tolerance of 0.020 Da and a parent ion tolerance of 10.0 PPM. Carbamidomethyl of cysteine was specified in Sequest and X! Tandem as a fixed modification. Deamidated of asparagine and glutamine and oxidation of methionine were specified in Sequest as variable modifications. Glu->pyro-Glu of the n-terminus, ammonia-loss of the n-terminus, gln->pyro-Glu of the n-terminus, deamidated of asparagine and glutamine and oxidation of methionine were specified in X! Tandem as variable modifications.

Scaffold (version Scaffold_4.7.2, Proteome Software Inc., Portland, OR) was used to validate MS/MS based peptide and protein identifications. Peptide identifications were accepted if they could be established at greater than 95.0% probability. Peptide Probabilities from X! Tandem were assigned by the Peptide Prophet algorithm (Keller et al., 2002) with Scaffold delta-mass correction. Peptide probabilities from Sequest were assigned by the Scaffold Local FDR algorithm. Protein identifications were accepted if they could be established at greater than 95.0% probability and contained at least 1 identified peptide. Protein probabilities were assigned by the Protein Prophet algorithm (Nesvizhskii et al., 2003). Proteins that contained similar peptides and could not be differentiated based on MS/MS analysis alone were grouped to satisfy the principles of parsimony. Proteins were annotated with GO terms from gene_association.goa_uniprot (downloaded Dec 14, 2015) (Ashburner et al., 2000). In addition, peak lists obtained from MS/MS spectra were identified independently using OMSSA version 2.1.9 (Geer et al., 2004), X!Tandem version X! Tandem Sledgehammer (2013.09.01.1) (Craig and Beavis, 2004), Andromeda version 1.5.3.4 (Cox et al., 2011), MS Amanda version 1.0.0.5242 (Dorfer et al., 2014), MS-GF+ version Beta (v10282) (Kim and Pevzner, 2014), Comet version 2015.02 rev. 3 (Eng et al., 2013), MyriMatch version 2.2.140 (Tabb et al., 2007) and Tide (Diament and Noble, 2011). The search was conducted using SearchGUI version 2.2.2 (Vaudel et al., 2011).

Protein identification was conducted against a concatenated target/decoy version (Elias and Gygi, 2010) of the Mus musculus (24797, >99.9%), Sus scrofa (1, <0.1%) complement of the UniProtKB (Apweiler et al., 2004) (version of December 2015, 24798, Mus Musculus) canonical and isoform sequences).The decoy sequences were created by reversing the target sequences in SearchGUI. The identification settings were as follows: Trypsin with a maximum of 2 missed cleavages; 10.0 ppm as MS1 and 0.5 Da as MS2 tolerances; fixed modifications: Carbamidomethylation of C (+57.021464 Da), variable modifications: Deamidation of N (+0.984016 Da), Deamidation of Q (+0.984016 Da), Oxidation of M (+15.994915 Da), Pyrolidone from E (--18.010565 Da), Pyrolidone from Q (--17.026549 Da), Pyrolidone from carbamidomethylated C (--17.026549 Da) and Acetylation of protein N-term (+42.010565 Da), fixed modifications during refinement procedure: Carbamidomethylation of C (+57.021464 Da).

Peptides and proteins were inferred from the spectrum identification results using PeptideShaker version 1.9.0 (Vaudel et al., 2015). Peptide Spectrum Matches (PSMs), peptides and proteins were validated at a 1.0% False Discovery Rate (FDR) estimated using the decoy-hit distribution. Spectrum counting abundance indexes were estimated using the Normalized Spectrum Abundance Factor (Powell et al., 2004) adapted for better handling of protein inference issues and peptide detectability. While the two independent protein algorithm searches largely matched with each other, a small subset of proteins were identified with high confidence using the SearchGUI/Peptideshaker platforms that were not identified with the ThermoFisher Scientific/Scaffold platforms.

The mass spectrometry data along with the identification results have been deposited to the ProteomeXchange Consortium (Vizcaíno et al., 2014) via the PRIDE partner repository (Martens et al., 2005) with the dataset identifier 466 PXD006046. During the review process this data can be accessed at https://www.ebi.ac.uk/pride/archive/ with the following information:

Username

Password: E0lS1QTw

### Dataset Filtering

Protein candidates from both pilot and primary LC/MS screens were subject to the following stringent criteria to build the KCC2 interactome. First pass filter grouping: at least 2 unique peptides and fold change of total spectra above 1.5. Second pass filter grouping: for proteins with only 1 unique peptide, consider whether (a) the protein isoform is an already validated KCC2 interactor in literature; (b) the protein isoform already appears in the first pass filter; (c) the protein isoform appears as a single-peptide interactor across experiments (using the same epitope KCC2 IPs / different epitope KCC2 IPs / different developmental time KCC2 IPs). If a particular protein isoform matches any of the above criteria, it gets shifted to the first pass filter grouping. Finally, the proteins that appear in the KCC2 interactome that are previously identified spurious interactors as identified in the CRAPome database (Mellacheruvu et al., 2013) were further eliminated. For the existing proteins a MaxP-SAINT score (Choi et al., 2012) was assigned and proteins were grouped as Gold, Silver or Bronze interactors prior to subsequent PPI (protein-protein interaction) network analysis. See **Figure 3 – Figure Supplement 3** for a detailed description of the path towards constructing the KCC2 interactome.

### Integrated PPI network analysis

Protein interactions were integrated with curated, high-throughput and predicted interactions from I2D ver. 2.3 database (Brown and Jurisica, 2007), FpClass high-confidence predictions (Kotlyar et al., 2014) and from the BioGRID database (Stark et al., 2006). Networks were visualized using Cytoscape ver. 3.3.0 (Shannon et al., 2003; Cline et al., 2007). Components of the KCC2 interactome were mapped to the excitatory synapse-enriched PSD proteome (Collins et al., 2006), or the inhibitory synapse-enriched GABA_A_R / GABA_B_R / NLGN2 / GlyR proteomes (Heller et al., 2012; Del Pino et al., 2014; Kang et al., 2014; Nakamura et al., 2016; Schwenk et al., 2016; Uezu et al., 2016).

### In vitro co-immunoprecipitation

HEK-293 and COS7 cells obtained from the ATCC were authenticated and checked for mycoplasma contamination. For co-immunoprecipitation experiments, cells were transfected with KCC2b-MYC, eGFP control, eGFP-PACSIN1/2/3, or eGFP-PACSIN1-deletion constructs (0.25 μg/construct) using Lipofectamine (Invitrogen) at 70% confluency. Thirty-six hours after transfection, cells were washed with ice-cold 1× PBS and lysed in modified RIPA buffer [50 mM Tris·HCl, pH 7.4, 150 mM NaCl, 1 mM EDTA, 1% Nonidet P-40, 0.1% SDS, 0.5% DOC, and protease inhibitors (Roche)]. Lysed cells were incubated on ice for 30 min and were centrifuged at 15,000 × g for 15 min at 4 °C. Cell lysates or solubilized membrane fractions (~0.2 – 0.5mg protein) were incubated with N-terminal KCC2b or anti-myc (CST #9B11, RRID AB_331783) antibodies on a rotating platform (4 h, 4 °C). Lysates were subsequently incubated with 20μl GammaBind IgG beads (GE Healthcare) on a rotating platform (1 h at 4 °C). After incubation, beads were washed twice with modified RIPA buffer, and twice with modified RIPA buffer minus detergents. Bound proteins were eluted with SDS sample buffer and subjected to SDS/PAGE along with 10% of input fraction and immunoblotted. **Figure 5e** is representative of 4 independent biological replicates; **Figure 5f** is representative of 3 independent biological replicates.

### BN-PAGE analysis and antibody-shift assay

Native-membrane fractions were prepared similarly as described (Swamy et al., 2006; Schwenk et al., 2012; Mahadevan et al., 2014, 2015). Antibody-shift assay and 2D BN-PAGE analysis of native-KCC2 complexes were performed as described previously (Mahadevan et al., 2014, 2015). Briefly, 50μg - 100μg of C_12_E_9_ solubilized complexes were pre-incubated for 1 hour with 10 μg of anti-N-terminal KCC2b antibody or chicken IgY whole molecule, prior to the addition of Coomassie blue G250. 1D-BN-PAGE was performed as described above using home-made 4% and 5% bis-tris gels as described (Swamy et al., 2006). After the completion of the gel run, excised BN-PAGE lanes were equilibrated in Laemmli buffer containing SDS and DTT for 15 minutes at room temperature to denature the native proteins. After a brief rinse in SDS-PAGE running buffer, the excised BN-PAGE lanes were placed on a 6% or 8% SDS-PAGE gel for separation in the second dimension. After standard electro-blotting of SDS-PAGE-resolved samples on nitrocellulose membrane, the blot was cut into two molecular weight ranges; the top blots were subjected to western blotting analysis with Rb anti-KCC2, and the bottom blots with Rb anti-PACSIN1. Antibody-shift experiments (**Figure 5d**) using hippocampal membranes are representative from 2 independent biological replicates.

### PACSIN overexpression and shRNA constructs

All PACSIN constructs used for overexpression and shRNA constructs have been previously validated for specificity (Anggono et al., 2013; Widagdo et al., 2016). The PACSIN1 shRNA-targeting sequence (sh#1, 5’-GCGCCAGCTCATCGAGAAA-3′) or control shRNA sequence was inserted into the pSuper vector system (Oligoengine) as described previously (Anggono et al., 2013). The efficiency and specificity of the PACSIN1 and control shRNA constructs were tested in HEK 293T cells overexpressing GFP-PACSIN1, and they were subsequently cloned into pAAV-U6 for lentiviral production (serotype AAV2/9).

### Hippocampal cultures and electrophysiology

Low-density cultures of dissociated mouse hippocampal neurons were prepared as previously described (Acton et al., 2012; Mahadevan et al., 2014). Experiments were performed after 10-13 days in culture (DIC). Electrophysiological recordings were performed using pipettes made from glass capillaries (WPI), as previously described (Acton et al., 2012; Mahadevan et al., 2014). For Cl^−^ loading experiments in whole-cell configuration, pipettes (5–7 MΩ) were filled with an internal solution containing: 90 mM K^+^-gluconate, 30 mM KCl, 10 mM HEPES, 0.2 mM EGTA, 4 mM ATP, 0.3 mM GTP, and 10 mM phosphocreatine (pH 7.4, 300 mOsm). For gramicidin perforated recordings, pipettes with a resistance of 7–12 MΩ were filled with an internal solution containing 150 mM KCl, 10 mM HEPES, and 50μg/ml gramicidin (pH 7.4, 300 mOsm). Cultured neurons were continuously perfused with standard extracellular solution. Cultured neurons were selected for electrophysiology based on the following criteria: (1) with a healthy oval or pyramidal-shaped cell body; (2) multiple clearly identifiable processes; (3) a cell body and proximal dendrites that were relatively isolated; (4) reporter fluorescence (if applicable). Recordings started when the series resistance dropped below 50 MΩ. IV-curves were made by depolarizing the membrane potential in steps, while simultaneously stimulating GABAergic transmission. A 20μM GABA puff was applied to the soma. A linear regression of the IPSC/P amplitude was used to calculate the voltage dependence of IPSC/Ps; the intercept of this line with the abscissa was taken as E_GABA_, and the slope of this line was taken as the synaptic conductance. The maximum current amplitude was taken as the largest absolute current recorded during the recordings performed for the E_GABA_ measurement. Electrophysiological values have not been corrected for the liquid junction potential of ~7 mV.

### Fixed immunostaining and confocal microscopy

DIV 12-14 cultured hippocampal neurons with were first rinsed with 1X PBS, and fixed in 4% paraformaldehyde for 10 min on ice followed by washing thrice with 1X PBS. Neurons were then permeabilized with 1X PBS containing 10% goat serum and 0.5% Triton X-100 for 30 minutes, followed by a 45 minute incubation with rabbit anti-KCC2 (Millipore 07-432) antibodies at 37 °C to detect endogenous proteins. Finally, neurons were washed thrice with 1X PBS and incubated with Alexa-fluor 555-conjugated goat anti-rabbit antibody for 45 minutes at 37°C. Neurons were imaged on a Leica TCS SP8 confocal system with a Leica DMI 6000 inverted microscope (Quorum Technologies). Cultured neurons were selected for immunostaining based on the following criteria: (1) with a healthy oval or pyramidal-shaped cell body; (2) multiple clearly identifiable processes; (3) a cell body and proximal dendrites that were relatively isolated; (4) reporter fluorescence (if applicable). Images were acquired using 3D Image Analysis software (Perkin Elmer). Images were obtained using a 63x 1.4-NA oil immersion objective. Imaging experiments were performed and analyzed in a blinded manner.

### Statistics

For electrophysiology and immunostaining data (**Figure 6**), ‘n’ values report the number of neurons, and were obtained from a minimum of three independent sets of cultured neurons (produced from different litters). Example recordings in **Figure 6a** are representative of n=9 (shRNA control) and n=11 (PACSIN1 shRNA). Example recordings in **Figure 6e** are representative of n=32 (shRNA control) and n=32 (PACSIN1 shRNA). Example recordings in **Figure 6f** are representative of n=23 (eGFP) and n=16 (PACSIN1-eGFP). Data in **Figure 6 b, c, e and f** are mean ± SEM. Statistical significance was determined using either SigmaStat or GraphPad Prism (version 5.01) software. Statistical significance in **Figure 6 b, c, d, e and f** was determined using Student’s t-tests (two-tailed); all data sets passed the normal distribution assumptions test. Statistical significance is noted as follows: * p < 0.05, ** p < 0.01, *** p < 0.001. Exact p and t values are reported in the Results text.

**Figure 1a, 5b, 5c, 5d** are representative of two independent biological replicates. **Figure 5f** is representative of three independent biological replicates. **Figure 5e** is representative of four independent biological replicates.

## ACKNOWLEDGEMENTS

We thank the following persons for technical assistance with LC/MS: Dr. Suzanne Ackloo at the Centre for Biological Timing and Cognition, University of Toronto; Drs. Paul Taylor and Jonathan Krieger, at the SPARC BioCentre, Hospital for Sick Children. We thank Drs. Harald Barsnes and Marc Vaudel, University of Bergen, for assistance with the PeptideShaker / CompOmics application. This study was funded by the following: Simons Foundation Autism Research Initiative to M.W. and Y.D.K,; CIHR training grant from the Sleep and Biological Rhythms Training Program, Toronto to V.M; The Academy of Finland grants to P.U. and M.S.A; The John T. Reid Charitable Trusts and Australian Research Council (ARC) project grant (DP170102402) to V.A.; CIHR foundation grant FRN-133431 to A.E; CIHR Operating Grant to M.W.

---

**Figure 1 – Figure Supplemental 1.**
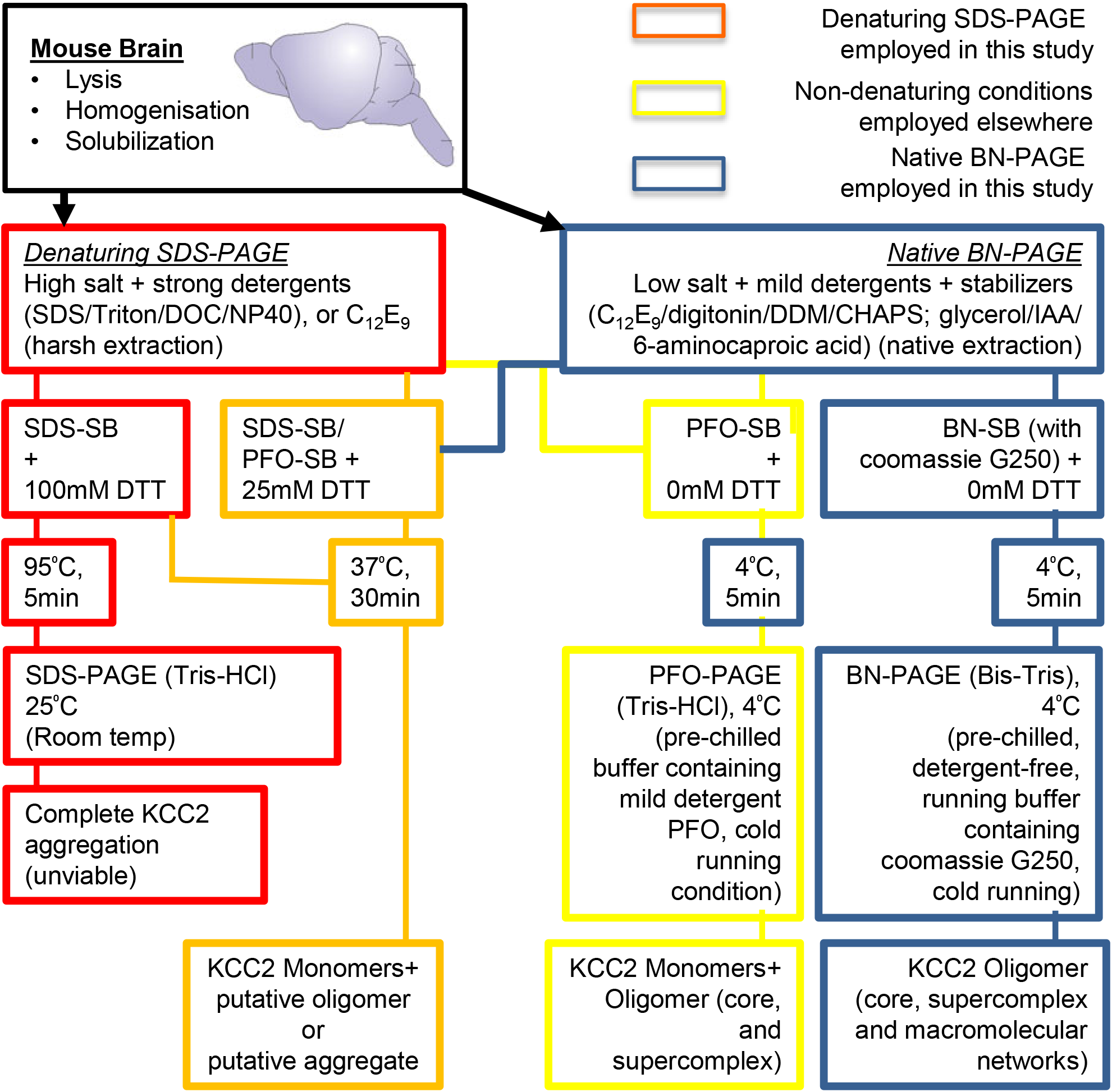
Workflow to enrich KCC2 complexes. SDS, sodium dodecyl sulfate; DOC, deoxycholate; NP40, Igepal-CA630; C_12_E_9_, nonaethylene glycol monododecyl ether; DDM, n-dodecyl-β-D-maltoside; PFO, perfluoro-octanoic acid; IAA, Iodoacetmide; BN, blue-native; SB, sample buffer for gel loading. RED/ORANGE lines and boxes indicate harsh KCC2 extraction conditions; YELLOW lines and boxes indicate intermediary KCC2 extraction conditions; BLUE lines and boxes indicate mild, native-KCC2 extraction conditions. The orange and yellow extraction/gel running strategies were employed for studying the stability of KCC2 oligomers (by subjecting them to harsh-to-mildly denaturing conditions). The blue extraction/gel running conditions were employed to study the composition of native-KCC2-oligomers.

**Figure 1 – Figure Supplemental 2.**
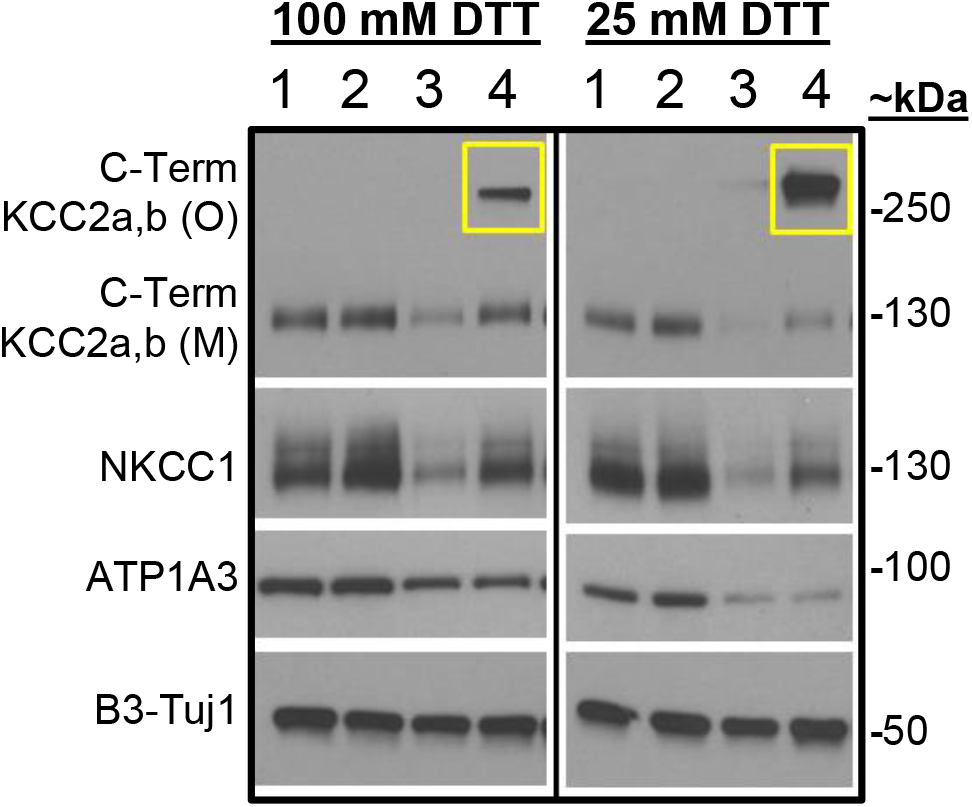
SDS-PAGE separation of solubilized membrane fractions. Obtained with high-salt Tris-HCl buffer containing the following detergents (<u>lane 1</u>: 1%Triton, 1%DOC; <u>lane 2</u>: 0.1%SDS, 0.5%DOC, 1%NP40 (RIPA); <u>lane 3</u>: 1%NP40 and lane 4: 1.5% C_12_E_9_). Samples were denatured in SDS-sample buffer containing 100mM DTT or 25mM DTT, @ 37°C for 30 min. Yellow-boxes indicate that C_12_E_9_-based native detergent enriches for more putative-KCC2 oligomers than the previously published KCC2 detergent extractions (lanes 1–3); and that the putative KCC2 oligomers are DTT-sensitive.

**Figure 2 – Figure Supplemental 1.**
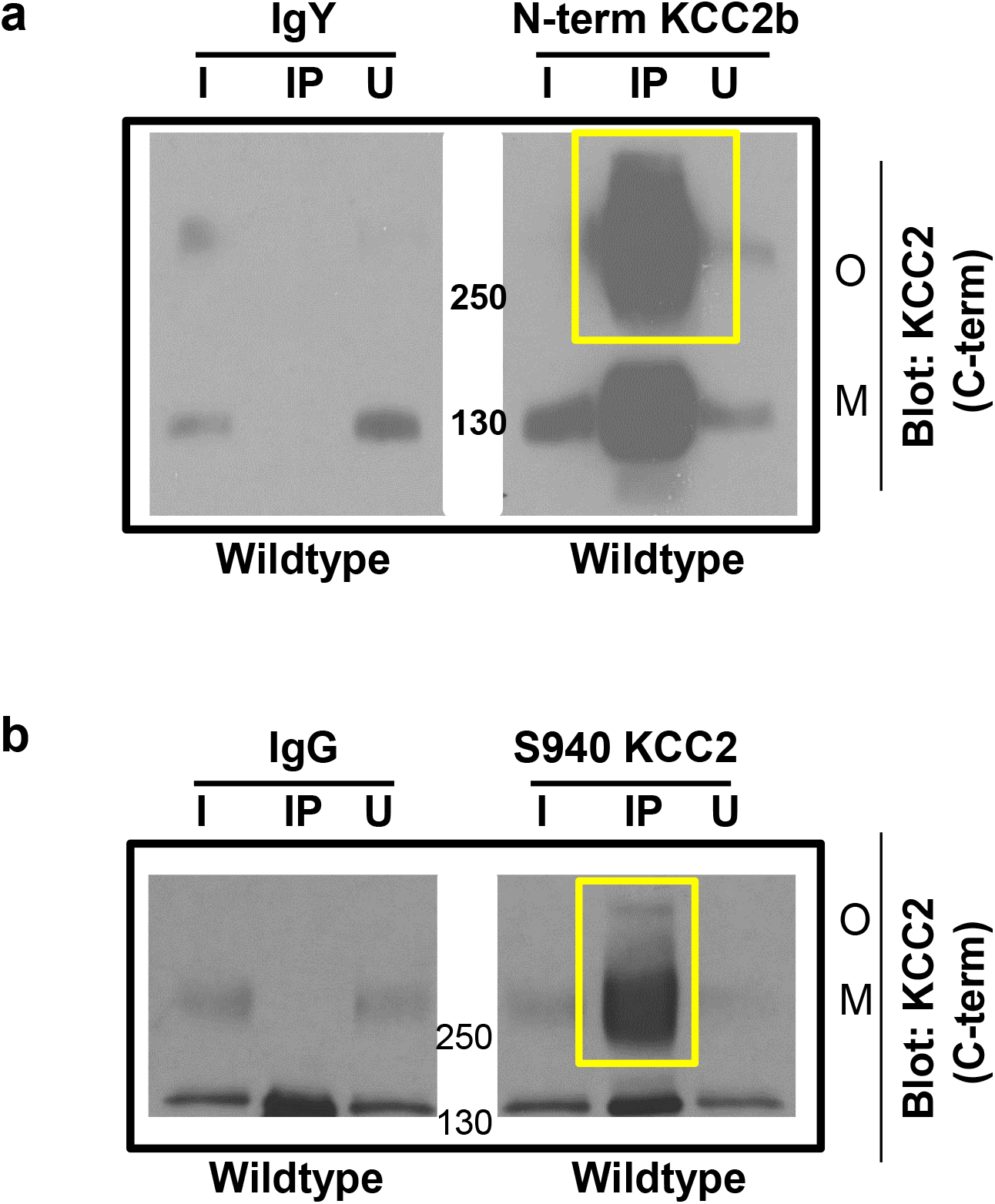
Validation of KCC2 antibodies for immunodepletion. Native KCC2 complexes from C_12_E_9_-solubilized whole-brain membrane fractions immunoprecipitated with pre-immune sera or anti-N-terminal KCC2b antibody **(a)** or anti-p^Ser^940 KCC2 antibody **(b)** and immunoblotted with the antibodies indicated at right (C-terminal KCC2 antibody). Representative example of 5 biological replicates. IP, immunoprecipitate; I, input fraction (1% of IP); U. unbound fraction (1%of IP); O. oligomer; M. monomer.

**Figure 3 – Figure Supplement 1.**
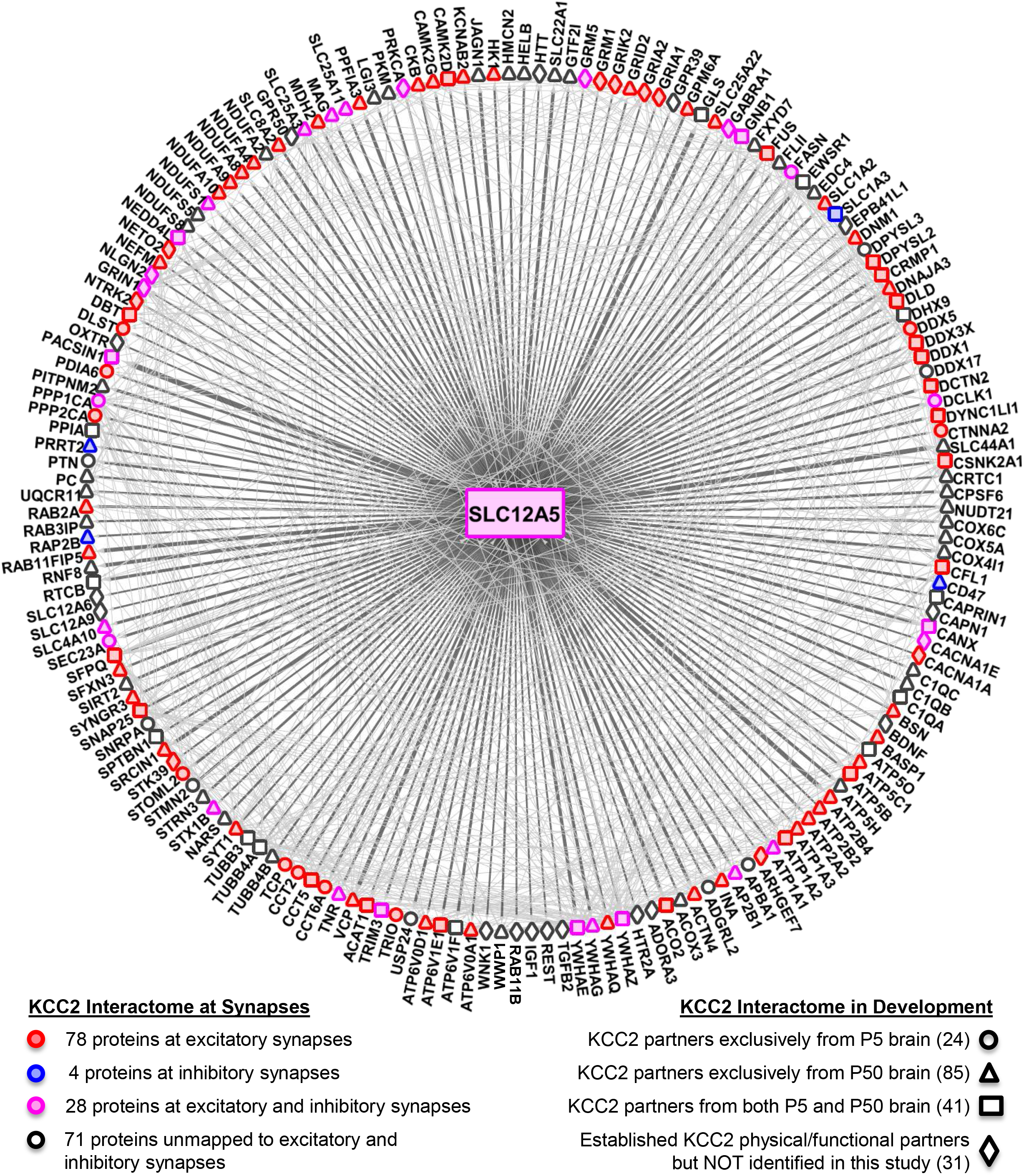
The SLC12A5/KCC2 interactome. The KCC2 interactome was mapped to the excitatory synapse-enriched PSD proteome, and the inhibitory synapse-enriched GABA_A_R / GABA_B_R / NLGN2 / GlyR proteomes. Circle/triangle/square-shaped nodes represent the KCC2 partners identified in this present study; diamond-shaped nodes represent the KCC2 partners not identified, but previously established as physical/functional partners of KCC2. Red/blue/pink-filled nodes represent synaptic-KCC2 partners; uncoloured nodes represent the putative-, non-synaptic KCC2 partners. The thickness of the edge represents the spectral enrichment (KCC2/IgG). See Supplemental Table 2 for the complete list of all proteins used for mapping.

**Figure 3 – Figure Supplement 2.**
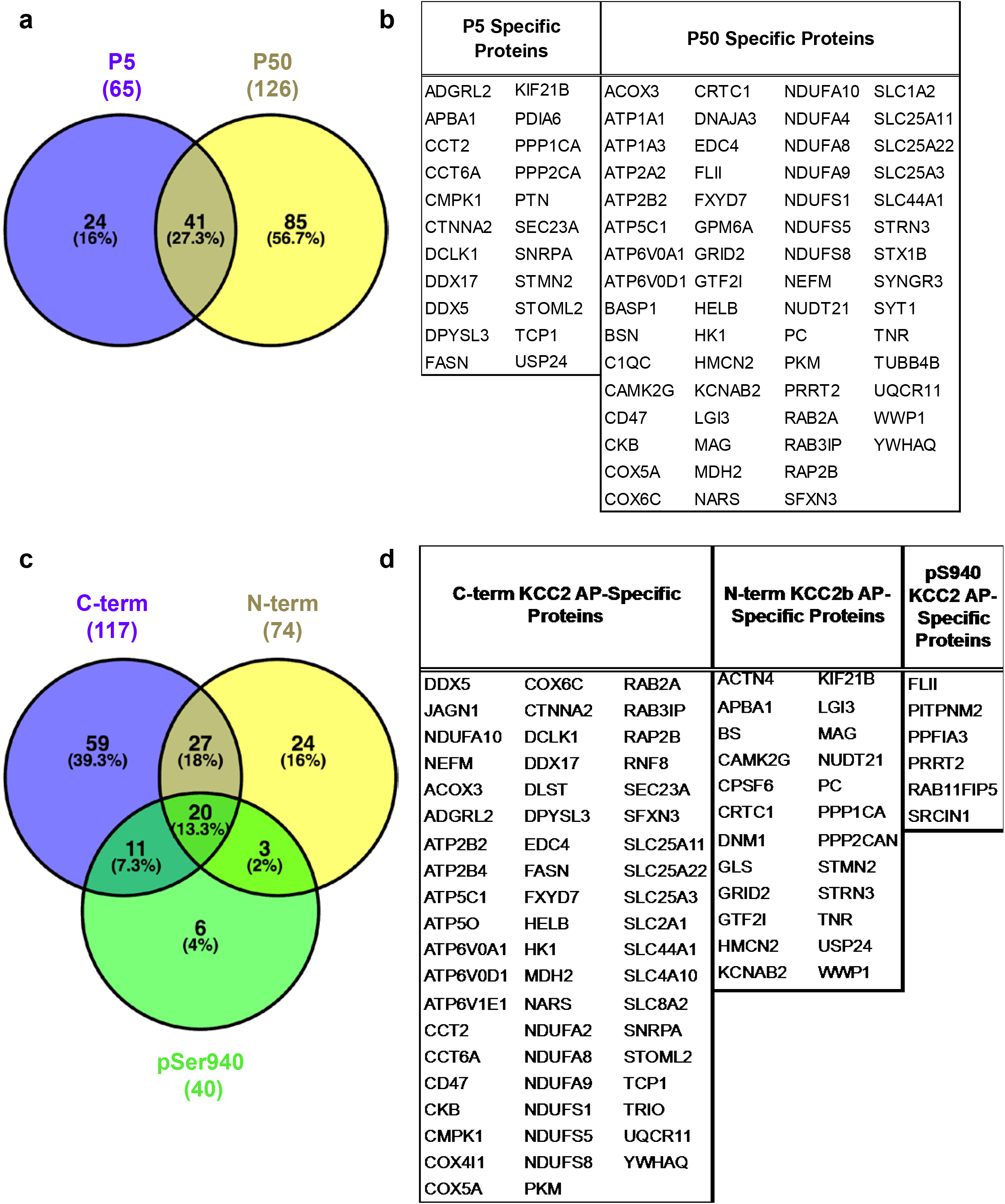
ME-AP proteomics identify the protein constituents of native KCC2. **(a)** Venn diagram comparison of the intersection of data obtained using N-term and C-term antibodies in developing and mature brain. **(b)** Proteins that appear exclusively with P5 or P50 KCC2-immunoprecipitates. **(c)** Similar to (a) but for data obtained using all three antibodies. **(d)** Proteins that appear exclusively to the individual KCC2 antibody.

**Figure 3 – Figure Supplement 3.**
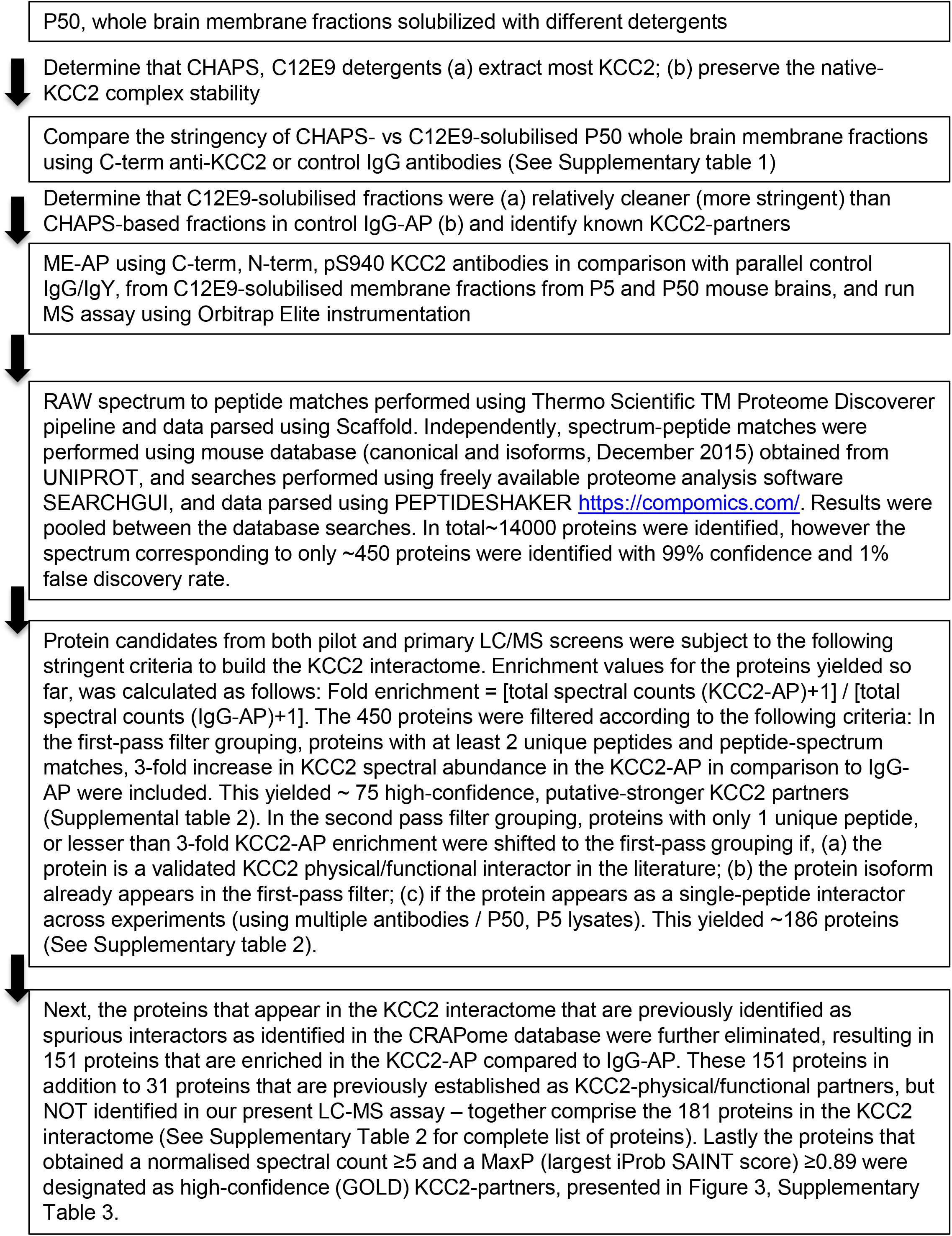
Workflow for curating the KCC2 interactome.

**Figure 5 – Figure Supplement 1.**
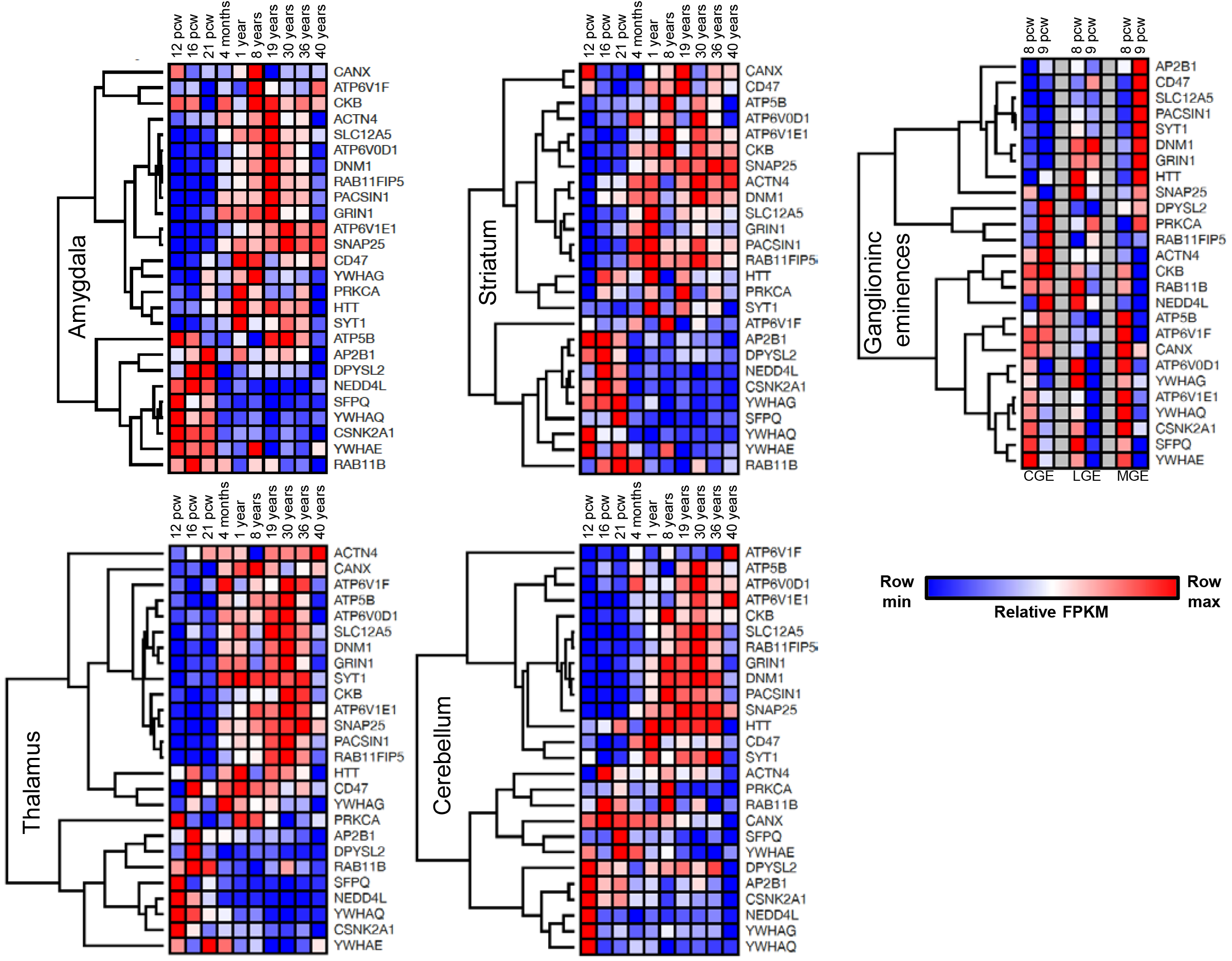
Spatiotemporal expression patterns of *SLC12A5* and members of receptor trafficking node of the KCC2 interactome in the human brain. The RNAseq data were analyzed for the above members across 5 brain regions including Amygdala, Striatum, Thalamus, Cerebellum and the ganglionic eminences at 8 different developmental periods.

**Figure 5 – Figure Supplement 2.**
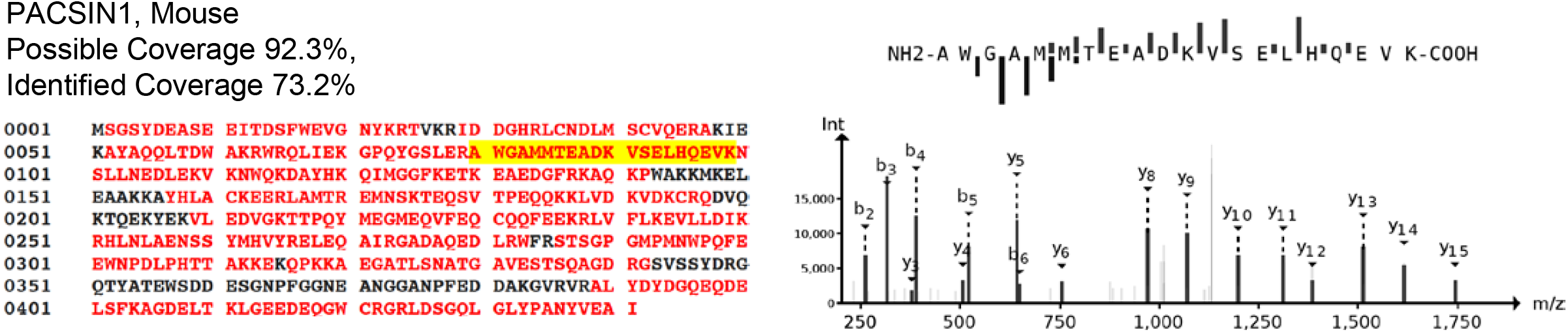
The primary amino acid sequence coverage of PACSIN1 (left), and protein coverage of PACSIN1 identified by MS analysis are indicated in red. MS/MS-spectra of a peptide unique for PACSIN1, highlighted in yellow. The MS/MS ion fragmentation of the corresponding amino acid sequence is indicated above the spectra (right).

## REFERENCES

Acton BA, Mahadevan V, Mercado A, Uvarov P, Ding Y, Pressey J, Airaksinen MS, Mount DB, Woodin M a. (2012) Hyperpolarizing GABAergic Transmission Requires the KCC2 C-Terminal ISO Domain. J Neurosci 32:8746–8751.

Anggono V, Koç-Schmitz Y, Widagdo J, Kormann J, Quan A, Chen C-M, Robinson PJ, Choi S-Y, Linden DJ, Plomann M, Huganir RL (2013) PICK1 interacts with PACSIN to regulate AMPA receptor internalization and cerebellar long-term depression. Proc Natl Acad Sci U S A 110:13976–13981

Anggono V, Smillie KJ, Graham ME, Valova VA, Cousin MA, Robinson PJ (2006) Syndapin I is the phosphorylation-regulated dynamin I partner in synaptic vesicle endocytosis. Nat Neurosci 9:752–760.

Apweiler R, Bairoch A, Wu CH, Barker WC, Boeckmann B, Ferro S, Gasteiger E, Huang H, Lopez R, Magrane M, Martin MJ, Natale DA, O’Donovan C, Redaschi N, Yeh L-SL (2004) UniProt: the Universal Protein knowledgebase. Nucleic Acids Res 32:D115–9.

Ashburner M et al. (2000) Gene ontology: tool for the unification of biology. The Gene Ontology Consortium. Nat Genet 5:25–29.

Babu M, Greenblatt JF, Emili A, Strynadka NCJ, Reithmeier RAF, Moraes TF (2010) Structure of a SLC26 anion transporter STAS domain in complex with acyl carrier protein: implications for E. coli YchM in fatty acid metabolism. Structure 18:1450–1462.

Bacaj T, Ahmad M, Jurado S, Malenka RC, Südhof TC (2015) Synaptic Function of Rab11Fip5: Selective Requirement for Hippocampal Long-Term Depression. J Neurosci 35:7460–7474

Banerjee A, Rikhye R V., Breton-Provencher V, Tang X, Li C, Li K, Runyan CA, Fu Z, Jaenisch R, Sur M (2016) Jointly reduced inhibition and excitation underlies circuit-wide changes in cortical processing in Rett syndrome. Proc Natl Acad Sci 113:E7287–E7296

Ben-Ari Y (2002) Excitatory actions of gaba during development: the nature of the nurture. Nat Rev Neurosci 3:728–739

Berkefeld H, Sailer CA, Bildl W, Rohde V, Thumfart J-O, Eble S, Klugbauer N, Reisinger E, Bischofberger J, Oliver D, Knaus H-G, Schulte U, Fakler B (2006) BKCa-Cav channel complexes mediate rapid and localized Ca2+-activated K+ signaling. Science 314:615–620.

Blaesse P, Airaksinen MS, Rivera C, Kaila K (2009) Cation-chloride cotransporters and neuronal function. Neuron 61:820–838

Blaesse P, Guillemin I, Schindler J, Schweizer M, Delpire E, Khiroug L, Friauf E, Nothwang HG (2006) Oligomerization of KCC2 correlates with development of inhibitory neurotransmission. J Neurosci 26:10407–10419

Blaesse P, Schmidt T (2014) K-Cl cotransporter KCC2-a moonlighting protein in excitatory and inhibitory synapse development and function. Pflugers Arch

Brown KR, Jurisica I (2007) Unequal evolutionary conservation of human protein interactions in interologous networks. Genome Biol 8:R95.

Carroll RC, Beattie EC, Xia H, Lüscher C, Altschuler Y, Nicoll RA, Malenka RC, von Zastrow M (1999) Dynamin-dependent endocytosis of ionotropic glutamate receptors. Proc Natl Acad Sci U S A 96:14112–14117.

Cellot G, Cherubini E (2014) GABAergic signaling as therapeutic target for autism spectrum disorders. Front Pediatr 2:70

Cembrowski MS, Wang L, Sugino K, Shields BC, Spruston N (2016) Hipposeq: a comprehensive RNA-seq database of gene expression in hippocampal principal neurons. Elife 5:e14997.

César-Razquin A, Snijder B, Frappier-Brinton T, Isserlin R, Gyimesi G, Bai X, Reithmeier RA, Hepworth D, Hediger MA, Edwards AM, Superti-Furga G (2015) A Call for Systematic Research on Solute Carriers. Cell 162:478–487

Chamma I, Chevy Q, Poncer JC, Lévi S (2012) Role of the neuronal K-Cl co-transporter KCC2 in inhibitory and excitatory neurotransmission. Front Cell Neurosci 6:5

Chamma I, Heubl M, Chevy Q, Renner M, Moutkine I, Eugène E, Poncer JC, Lévi S (2013) Activity-dependent regulation of the K/Cl transporter KCC2 membrane diffusion, clustering, and function in hippocampal neurons. J Neurosci 33:15488–15503.

Chevy Q, Heubl M, Goutierre M, Backer S, Moutkine I, Eugène E, Bloch-Gallego E, Lévi S, Poncer JC (2015) KCC2 Gates Activity-Driven AMPA Receptor Traffic through Cofilin Phosphorylation. J Neurosci 35:15772–15786

Chiu CQ, Lur G, Morse TM, Carnevale NT, Ellis-Davies GCR, Higley MJ (2013) Compartmentalization of GABAergic inhibition by dendritic spines. Science 340:759–762

Choi H, Liu G, Mellacheruvu D, Tyers M, Gingras A-C, Nesvizhskii AI (2012) Analyzing protein-protein interactions from affinity purification-mass spectrometry data with SAINT. Curr Protoc Bioinforma Chapter 8:Unit8.15.

Clayton EL, Anggono V, Smillie KJ, Chau N, Robinson PJ, Cousin MA (2009) The Phospho-Dependent Dynamin-Syndapin Interaction Triggers Activity-Dependent Bulk Endocytosis of Synaptic Vesicles. J Neurosci 29:7706–7717.

Cline MS et al. (2007) Integration of biological networks and gene expression data using Cytoscape. Nat Protoc 2:2366–2382.

Collins MO, Husi H, Yu L, Brandon JM, Anderson CNG, Blackstock WP, Choudhary JS, Grant SGN (2006) Molecular characterization and comparison of the components and multiprotein complexes in the postsynaptic proteome. J Neurochem 97 Suppl 1:16–23

Coull JAM, Boudreau D, Bachand K, Prescott S a, Nault F, Sík A, De Koninck P, De Koninck Y (2003) Trans-synaptic shift in anion gradient in spinal lamina I neurons as a mechanism of neuropathic pain. Nature 424:938–942.

Cox J, Neuhauser N, Michalski A, Scheltema RA, Olsen J V, Mann M (2011) Andromeda: a peptide search engine integrated into the MaxQuant environment. J Proteome Res 10:1794–1805.

Craig R, Beavis RC (2004) TANDEM: matching proteins with tandem mass spectra. Bioinformatics 20:1466–1467.

Del Pino I, Koch D, Schemm R, Qualmann B, Betz H, Paarmann I (2014) Proteomic analysis of glycine receptor β subunit (GlyRβ)-interacting proteins: evidence for syndapin I regulating synaptic glycine receptors. J Biol Chem 289:11396–11409.

Diament BJ, Noble WS (2011) Faster SEQUEST searching for peptide identification from tandem mass spectra. J Proteome Res 10:3871–3879.

Dorfer V, Pichler P, Stranzl T, Stadlmann J, Taus T, Winkler S, Mechtler K (2014) MS Amanda, a universal identification algorithm optimized for high accuracy tandem mass spectra. J Proteome Res 13:3679–3684.

Doyon N, Vinay L, Prescott SA, De Koninck Y (2016) Chloride Regulation: A Dynamic Equilibrium Crucial for Synaptic Inhibition. Neuron 89:1157–1172.

Elias JE, Gygi SP (2010) Target-decoy search strategy for mass spectrometry-based proteomics. Methods Mol Biol 604:55–71.

Eng JK, Jahan TA, Hoopmann MR (2013) Comet: an open-source MS/MS sequence database search tool. Proteomics 13:22–24.

Fiumelli H, Briner A, Puskarjov M, Blaesse P, Belem BJ, Dayer AG, Kaila K, Martin JL, Vutskits L (2013) An ion transport-independent role for the cation-chloride cotransporter kcc2 in dendritic spinogenesis in vivo. Cereb Cortex 23:378–388.

Ford A, Castonguay XA, Cottet XM, Little JW, Chen Z, Symons-liguori XAM, Doyle T, Egan TM, Vanderah TW, Konnick Y De, Tosh DK, Jacobson XK a, Salvemini D (2015) Engagement of the GABA to KCC2 Signaling Pathway Contributes to the Analgesic Effects of A3AR Agonists in Neuropathic Pain. J Neurosci 35:6057–6067.

Friedel P, Kahle KT, Zhang J, Hertz N, Pisella LI, Buhler E, Schaller F, Duan J, Khanna AR, Bishop PN, Shokat KM, Medina I (2015) WNK1-regulated inhibitory phosphorylation of the KCC2 cotransporter maintains the depolarizing action of GABA in immature neurons. Sci Signal 8:ra65.

Garbarini N, Delpire E (2008) The RCC1 domain of protein associated with Myc (PAM) interacts with and regulates KCC2. Cell Physiol Biochem 22:31–44

Gauvain G, Chamma I, Chevy Q, Cabezas C, Irinopoulou T, Bodrug N, Carnaud M, Lévi S, Poncer JC (2011) The neuronal K-Cl cotransporter KCC2 influences postsynaptic AMPA receptor content and lateral diffusion in dendritic spines. Proc Natl Acad Sci U S A 108:15474–15479

Geer LY, Markey SP, Kowalak JA, Wagner L, Xu M, Maynard DM, Yang X, Shi W, Bryant SH (2004) Open mass spectrometry search algorithm. J Proteome Res 3:958–964.

Gulyas AI, Sik A, Payne JA, Kaila K, Freund TF, Gulya AI (2001) The KCl cotransporter, KCC2, is highly expressed in the vicinity of excitatory synapses in the rat hippocampus. Eur J Neurosci 13:2205–2217

Heller EA, Zhang W, Selimi F, Earnheart JC, Ślimak MA, Santos-Torres J, Ibañez-Tallon I, Aoki C, Chait BT, Heintz N (2012) The biochemical anatomy of cortical inhibitory synapses Nitabach MN, ed. PLoS One 7:e39572.

Higley MJ (2014) Localized GABAergic inhibition of dendritic Ca2+ signalling. Nat Rev Neurosci 15:567–572

Horn Z, Ringstedt T, Blaesse P, Kaila K, Herlenius E (2010) Premature expression of KCC2 in embryonic mice perturbs neural development by an ion transport-independent mechanism. Eur J Neurosci 31:2142–2155

Huberfeld G, Wittner L, Clemenceau S, Baulac M, Kaila K, Miles R, Rivera C (2007) Perturbed chloride homeostasis and GABAergic signaling in human temporal lobe epilepsy. J Neurosci 27:9866–9873

Husi H, Ward MA, Choudhary JS, Blackstock WP, Grant SG (2000) Proteomic analysis of NMDA receptor-adhesion protein signaling complexes. Nat Neurosci 3:661–669

Ikeda K, Onimaru H, Yamada J, Inoue K, Ueno S, Onaka T, Toyoda H, Arata A, Ishikawa T, Taketo MM, Fukuda A, Kawakami K (2004) Malfunction of respiratory-related neuronal activity in Na+, K+-ATPase alpha2 subunit-deficient mice is attributable to abnormal Cl-homeostasis in brainstem neurons. J Neurosci 24:10693–10701

Inoue K, Ueno S, Fukuda A (2004) Interaction of neuron-specific K+-Cl-cotransporter, KCC2, with brain-type creatine kinase. FEBS Lett 564:131–135.

Inoue K, Yamada J, Ueno S, Fukuda A (2006) Brain-type creatine kinase activates neuron-specific K+-Cl-co-transporter KCC2. J Neurochem 96:598–608.

Ivakine EA, Acton BA, Mahadevan V, Ormond J, Tang M, Pressey JC, Huang MY, Ng D, Delpire E, Salter MW, Woodin MA, McInnes RR (2013) Neto2 is a KCC2 interacting protein required for neuronal Cl-regulation in hippocampal neurons. Proc Natl Acad Sci U S A 110:3561–3566

Kahle KKT, Deeb TTZ, Puskarjov M, Silayeva L, Liang B, Kaila K, Moss SJ (2013) Modulation of neuronal activity by phosphorylation of the K-Cl cotransporter KCC2. Trends Neurosci 36:726–737.

Kahle KT et al. (2014) Genetically encoded impairment of neuronal KCC2 cotransporter function in human idiopathic generalized epilepsy. EMBO Rep 15:766–774

Kahle KT, Staley KJ, Nahed B V, Gamba G, Hebert SC, Lifton RP, Mount DB (2008) Roles of the cation-chloride cotransporters in neurological disease. Nat Clin Pract Neurol 4:490–503

Kaila K, Price T, Payne J, Puskarjov M, Voipio J (2014) Cation-chloride cotransporters in neuronal development, plasticity and disease. Nat Rev Neurosci 15:637–654

Kang Y, Ge Y, Cassidy RM, Lam V, Luo L, Moon K-M, Lewis R, Molday RS, Wong ROL, Foster LJ, Craig AM (2014) A combined transgenic proteomic analysis and regulated trafficking of neuroligin-2. J Biol Chem 289:29350–29364.

Keller A, Nesvizhskii A, Kolker E, Aebersold R (2002) Empirical statistical model to estimate the accuracy of peptide identifications made by MS/MS and database search. Anal Chem 74:5383–5392.

Kessels MM, Qualmann B (2004) The syndapin protein family: linking membrane trafficking with the cytoskeleton. J Cell Sci 117:3077–3086.

Kessels MM, Qualmann B (2006) Syndapin Oligomers Interconnect the Machineries for Endocytic Vesicle Formation and Actin Polymerization. J Biol Chem 281:13285–13299.

Kessels MM, Qualmann B (2015) Different functional modes of BAR domain proteins in formation and plasticity of mammalian postsynapses. J Cell Sci:1–9.

Kim S, Pevzner PA (2014) MS-GF+ makes progress towards a universal database search tool for proteomics. Nat Commun 5:5277.

Kotlyar M, Pastrello C, Pivetta F, Lo Sardo A, Cumbaa C, Li H, Naranian T, Niu Y, Ding Z, Vafaee F, Broackes-Carter F, Petschnigg J, Mills GB, Jurisicova A, Stagljar I, Maestro R, Jurisica I (2014) In silico prediction of physical protein interactions and characterization of interactome orphans. Nat Methods 12:79–84

Lee HHC, Deeb TZ, Walker JA, Davies PA, Moss SJ (2011) NMDA receptor activity downregulates KCC2 resulting in depolarizing GABAA receptor-mediated currents. Nat Neurosci 14:736–743

Lee HHC, Walker JA, Williams JR, Goodier RJ, Payne JA, Moss SJ (2007) Direct protein kinase C-dependent phosphorylation regulates the cell surface stability and activity of the potassium chloride cotransporter KCC2. J Biol Chem 282:29777–29784

Lee SH, Liu L, Wang YT, Sheng M (2002) Clathrin adaptor AP2 and NSF interact with overlapping sites of GluR2 and play distinct roles in AMPA receptor trafficking and hippocampal LTD. Neuron 36:661–674.

Leonzino M, Busnelli M, Antonucci F, Verderio C, Mazzanti M, Chini B (2016) The Timing of the Excitatory-to-Inhibitory GABA Switch Is Regulated by the Oxytocin Receptor via KCC2. Cell Rep 15.

Li H, Khirug S, Cai C, Ludwig A, Blaesse P, Kolikova J, Afzalov R, Coleman SK, Lauri S, Airaksinen MS, Keinänen K, Khiroug L, Saarma M, Kaila K, Rivera C (2007) KCC2 interacts with the dendritic cytoskeleton to promote spine development. Neuron 56:1019–1033

Lin L, Yee SW, Kim RB, Giacomini KM (2015) SLC transporters as therapeutic targets: emerging opportunities. Nat Rev Drug Discov 14:543–560

Llano O, Smirnov S, Soni S, Golubtsov A, Guillemin I, Hotulainen P, Medina I, Nothwang HG, Rivera C, Ludwig A, 1Neuroscience (2015) KCC2 regulates actin dynamics in spines via interaction with betaPIX. J Cell Biol 209:671–686.

Mahadevan V, Dargaei Z, Ivakine EA, Hartmann A-M, Ng D, Chevrier J, Ormond J, Nothwang HG, McInnes RR, Woodin MA (2015) Neto2-null mice have impaired GABAergic inhibition and are susceptible to seizures. Front Cell Neurosci 9:368

Mahadevan V, Pressey JC, Acton BA, Uvarov P, Huang MY, Chevrier J, Puchalski A, Li CM, Ivakine EA, Airaksinen MS, Delpire E, McInnes RR, Woodin MA (2014) Kainate Receptors Coexist in a Functional Complex with KCC2 and Regulate Chloride Homeostasis in Hippocampal Neurons. Cell Rep 7:1762–1770

Mahadevan V, Woodin MA (2016) Regulation of neuronal chloride homeostasis by neuromodulators. J Physiol

Markkanen M, Karhunen T, Llano O, Ludwig A, Rivera C, Uvarov P, Airaksinen MS (2014) Distribution of neuronal KCC2a and KCC2b isoforms in mouse CNS. J Comp Neurol 522:1897–1914.

Martens L, Hermjakob H, Jones P, Adamski M, Taylor C, States D, Gevaert K, Vandekerckhove J, Apweiler R (2005) PRIDE: the proteomics identifications database. Proteomics 5:3537–3545.

Medina I, Friedel P, Rivera C, Kahle KT, Kourdougli N, Uvarov P, Pellegrino C (2014) Current view on the functional regulation of the neuronal K(+)-Cl(−) cotransporter KCC2. Front Cell Neurosci 8:27

Mellacheruvu D et al. (2013) The CRAPome: a contaminant repository for affinity purification-mass spectrometry data. Nat Methods 10:730–736.

Modregger J, Ritter B, Witter B, Paulsson M, Plomann M (2000) All three PACSIN isoforms bind to endocytic proteins and inhibit endocytosis. J Cell Sci 113:4511–4521.

Müller CS, Haupt A, Bildl W, Schindler J, Knaus H-G, Meissner M, Rammner B, Striessnig J, Flockerzi V, Fakler B, Schulte U (2010) Quantitative proteomics of the Cav2 channel nano-environments in the mammalian brain. Proc Natl Acad Sci U S A 107:14950–14957.

Nakamura Y, Morrow DH, Modgil A, Huyghe D, Deeb TZ, Lumb MJ, Davies PA, Moss SJ (2016) Proteomic characterization of inhibitory synapses using a novel pHluorin-tagged GABAAR α2 subunit knock-in mouse. J Biol Chem 291:12394–12407.

Nelson SB, Valakh V (2015) Excitatory/Inhibitory Balance and Circuit Homeostasis in Autism Spectrum Disorders. Neuron 87:684–698.

Nesvizhskii A, Keller A, Kolker E, Aebersold R (2003) A statistical model for identifying proteins by tandem mass spectrometry. Anal Chem 75:4646–4658.

Nesvizhskii AI, Aebersold R (2005) Interpretation of shotgun proteomic data: the protein inference problem. Mol Cell Proteomics 4:1419–1440.

Pérez-Otaño I, Luján R, Tavalin SJ, Plomann M, Modregger J, Liu X-B, Jones EG, Heinemann SF, Lo DC, Ehlers MD (2006) Endocytosis and synaptic removal of NR3A-containing NMDA receptors by PACSIN1/syndapin1. Nat Neurosci 9:611–621

Pin J-P, Bettler B (2016) Organization and functions of mGlu and GABAB receptor complexes. Nature 540:60–68.

Powell DW, Weaver CM, Jennings JL, McAfee KJ, He Y, Weil PA, Link AJ (2004) Cluster analysis of mass spectrometry data reveals a novel component of SAGA. Mol Cell Biol 24:7249–7259.

Pressey J, Mahadevan V, Khademullah C, Dargaei Z, Chevrier J, Ye W, Huang M, Chauhan A, Meas S, Uvarov P, Airaksinen M, Woodin M (2017) A Kainate Receptor Subunit Promotes the Recycling of the Neuron-Specific K+-Cl-Cotransporter KCC2 in Hippocampal Neurons. J Biol Chem 292:6190–6201.

Puskarjov M, Ahmad F, Kaila K, Blaesse P (2012) Activity-Dependent Cleavage of the K-Cl Cotransporter KCC2 Mediated by Calcium-Activated Protease Calpain. J Neurosci 32:11356–11364.

Puskarjov M, Seja P, Heron SE, Williams TC, Ahmad F, Iona X, Oliver KL, Grinton BE, Vutskits L, Scheffer IE, Petrou S, Blaesse P, Dibbens LM, Berkovic SF, Kaila K (2014) A variant of KCC2 from patients with febrile seizures impairs neuronal Cl-extrusion and dendritic spine formation. EMBO Rep 15:1–7.

Ramachandran N et al. (2013) VMA21 deficiency prevents vacuolar ATPase assembly and causes autophagic vacuolar myopathy. Acta Neuropathol 125:439–457.

Rivera C, Voipio J, Payne JA, Ruusuvuori E, Lahtinen H, Lamsa K, Pirvola U, Saarma M, Kaila K (1999) The K+/Cl-co-transporter KCC2 renders GABA hyperpolarizing during neuronal maturation. Nature 397:251–255

Romero F (2009) Solubilization and partial characterization of ouabain-insensitive Na(+)-ATPase from basolateral plasma membranes of the intestinal epithelial cells. Investig clínica 50:303–314.

Roussa E, Speer JM, Chudotvorova I, Khakipoor S, Smirnov S, Rivera C, Krieglstein K (2016) K+/Cl− cotransporter KCC2 membrane trafficking and functionality is regulated by transforming growth factor beta 2. J Cell Sci 129:3485–3498.

Saitsu H, Watanabe M, Akita T, Ohba C, Sugai K, Ong WP, Shiraishi H, Yuasa S, Matsumoto H, Beng KT, Saitoh S, Miyatake S, Nakashima M, Miyake N, Kato M, Fukuda A, Matsumoto N (2016) Impaired neuronal KCC2 function by biallelic SLC12A5 mutations in migrating focal seizures and severe developmental delay. Sci Rep 6:30072.

Sanz-Clemente A, Matta JA, Isaac JTR, Roche KW (2010) Casein kinase 2 regulates the NR2 subunit composition of synaptic NMDA receptors. Neuron 67:984–996.

Sarkar J, Wakefield S, MacKenzie G, Moss SJ, Maguire J (2011) Neurosteroidogenesis Is Required for the Physiological Response to Stress: Role of Neurosteroid-Sensitive GABAA Receptors. J Neurosci 31:18198–18210.

Schulte U, Müller CS, Fakler B (2011) Ion channels and their molecular environments-Glimpses and insights from functional proteomics. Semin Cell Dev Biol 22:132–144.

Schwenk J, Baehrens D, Haupt A, Bildl W, Boudkkazi S, Roeper J, Fakler B, Schulte U (2014) Regional Diversity and Developmental Dynamics of the AMPA-Receptor Proteome in the Mammalian Brain. Neuron 84:41–54

Schwenk J, Harmel N, Brechet A, Zolles G, Berkefeld H, Müller CS, Bildl W, Baehrens D, Hüber B, Kulik A, Klöcker N, Schulte U, Fakler B (2012) High-resolution proteomics unravel architecture and molecular diversity of native AMPA receptor complexes. Neuron 74:621–633

Schwenk J, Metz M, Zolles G, Turecek R, Fritzius T, Bildl W, Tarusawa E, Kulik A, Unger A, Ivankova K, Seddik R, Tiao JY, Rajalu M, Trojanova J, Rohde V, Gassmann M, Schulte U, Fakler B, Bettler B (2010) Native GABA(B) receptors are heteromultimers with a family of auxiliary subunits. Nature 465:231–235

Schwenk J, Pérez-garci E, Schneider A, Kollewe A, Gauthier-kemper A, Fritzius T, Raveh A, Dinamarca MC, Hanuschkin A, Bildl W, Klingauf J, Gassmann M (2016) Modular composition and dynamics of native GABA B receptor complexes identified by high-resolution proteomics. Nat Neurosci 19:233–242.

Selak S, Paternain A V., Aller IM, Picó E, Rivera R, Lerma J (2009) A Role for SNAP25 in Internalization of Kainate Receptors and Synaptic Plasticity. Neuron 63:357–371.

Shannon P, Markiel A, Ozier O, Baliga NS, Wang JT, Ramage D, Amin N, Schwikowski B, Ideker T (2003) Cytoscape: a software environment for integrated models of biomolecular interaction networks. Genome Res 13:2498–2504.

Silayeva L, Deeb TZ, Hines RM, Kelley MR, Munoz MB, Lee HHC, Brandon NJ, Dunlop J, Maguire J, Davies PA, Moss SJ (2015) KCC2 activity is critical in limiting the onset and severity of status epilepticus. Proc Natl Acad Sci U S A 112:3523–3528

Snijder B, Frappier-brinton T, Isserlin R, Gyimesi G, Bai X, Reithmeier RA, Hepworth D, Hediger MA, Edwards AM, Superti-furga G (2015) Perspective A Call for Systematic Research on Solute Carriers. Cell 162:478–487.

Stark C, Breitkreutz B-J, Reguly T, Boucher L, Breitkreutz A, Tyers M (2006) BioGRID: a general repository for interaction datasets. Nucleic Acids Res 34:D535–9.

Stödberg T et al. (2015) Mutations in SLC12A5 in epilepsy of infancy with migrating focal seizures. Nat Commun 6:8038.

Swamy M, Siegers GM, Minguet S, Wollscheid B, Schamel WWA (2006) Blue native polyacrylamide gel electrophoresis (BN-PAGE) for the identification and analysis of multiprotein complexes. Sci STKE 2006:p14

Tabb DL, Fernando CG, Chambers MC (2007) MyriMatch: highly accurate tandem mass spectral peptide identification by multivariate hypergeometric analysis. J Proteome Res 6:654–661.

Tang X, Kim J, Zhou L, Wengert E, Zhang L, Wu Z, Carromeu C, Muotri AR (2015) KCC2 rescues functional deficits in human neurons derived from patients with Rett syndrome. Proc Natl Acad Sci 113:751–756.

Tang X, Kim J, Zhou L, Wengert E, Zhang L, Wu Z, Carromeu C, Muotri AR, Marchetto MCN, Gage FH, Chen G (2016) KCC2 rescues functional deficits in human neurons derived from patients with Rett syndrome. Proc Natl Acad Sci U S A 113:751–756.

Tao R, Li C, Newburn EN, Ye T, Lipska BK, Herman MM, Weinberger DR, Kleinman JE, Hyde TM (2012) Transcript-specific associations of SLC12A5 (KCC2) in human prefrontal cortex with development, schizophrenia, and affective disorders. J Neurosci 32:5216–5222

Toda T, Ishida K, Kiyama H, Yamashita T, Lee S (2014) Down-Regulation of KCC2 Expression and Phosphorylation in Motoneurons, and Increases the Number of in Primary Afferent Projections to Motoneurons in Mice with Post-Stroke Spasticity. PLoS One 9:e114328

Uezu A, Kanak DJ, Bradshaw TWA, Soderblom EJ, Catavero CM, Burette AC, Weinberg RJ, Soderling SH (2016) Identification of an elaborate complex mediating postsynaptic inhibition. Science (80-) 353:1123–1129.

Uvarov P, Ludwig A, Markkanen M, Pruunsild P, Kaila K, Delpire E, Timmusk T, Rivera C, Airaksinen MS (2007) A novel N-terminal isoform of the neuron-specific K-Cl cotransporter KCC2. J Biol Chem 282:30570–30576

Uvarov P, Ludwig A, Markkanen M, Soni S, Hübner CA, Rivera C, Airaksinen MS (2009) Coexpression and heteromerization of two neuronal K-Cl cotransporter isoforms in neonatal brain. J Biol Chem 284:13696–13704

Vandenberghe W, Nicoll RA, Bredt DS (2005) Stargazin is an AMPA receptor auxiliary subunit. Proc Natl Acad Sci U S A 102:485–490.

Vaudel M, Barsnes H, Berven FS, Sickmann A, Martens L (2011) SearchGUI: An open-source graphical user interface for simultaneous OMSSA and X!Tandem searches. Proteomics 11:996–999.

Vaudel M, Burkhart JM, Zahedi RP, Oveland E, Berven FS, Sickmann A, Martens L, Barsnes H (2015) PeptideShaker enables reanalysis of MS-derived proteomics data sets. Nat Biotechnol 33:22–24.

Vizcaíno JA et al. (2014) ProteomeXchange provides globally coordinated proteomics data submission and dissemination. Nat Biotechnol 32:223–226.

Wake H, Watanabe M, Moorhouse AJ, Kanematsu T, Horibe S, Matsukawa N, Asai K, Ojika K, Hirata M, Nabekura J (2007) Early changes in KCC2 phosphorylation in response to neuronal stress result in functional downregulation. J Neurosci 27:1642–1650

Watanabe M, Wake H, Moorhouse AJ, Nabekura J (2009) Clustering of neuronal K+-Cl− cotransporters in lipid rafts by tyrosine phosphorylation. J Biol Chem 284:27980–27988

Wenz M, Hartmann AM, Friauf E, Nothwang HG (2009) CIP1 is an activator of the K+-Cl− cotransporter KCC2. Biochem Biophys Res Commun 381:388–392

Widagdo J, Fang H, Jang SE, Anggono V (2016) PACSIN1 regulates the dynamics of AMPA receptor trafficking. Sci Rep 6:31070.

Williams JR, Sharp JW, Kumari VG, Wilson M, Payne JA (1999) The Neuron-specific K-Cl Cotransporter, KCC2. J Biol Chem 274:12656–12664.

Woo N-S, Lu J, England R, McClellan R, Dufour S, Mount DB, Deutch AY, Lovinger DM, Delpire E (2002) Hyperexcitability and epilepsy associated with disruption of the mouse neuronal-specific K-Cl cotransporter gene. Hippocampus 12:258–268

